# T cell Egress via Lymphatic Vessels Limits the Intratumoral T cell Repertoire in Melanoma

**DOI:** 10.1101/2022.05.30.494080

**Authors:** Maria M. Steele, Ian D. Dryg, Dhaarini Murugan, Julia Femel, Haley du Bois, Cameron Hill, Sancy A. Leachman, Young H. Chang, Lisa M. Coussens, Amanda W. Lund

## Abstract

Antigen-specific CD8^+^ T cell accumulation in tumors is a prerequisite for effective immunotherapy, and yet, the mechanisms of lymphocyte transit remain poorly defined. We find that tumor-associated lymphatic vessels control T cell exit from tumors via the chemokine CXCL12, and intratumoral antigen encounter tunes CXCR4 expression on effector CD8^+^ T cells. Only high affinity antigen downregulates CXCR4 and upregulates the CXCL12 decoy receptor, ACKR3, thereby reducing CXCL12 sensitivity and promoting T cell retention. A diverse repertoire of functional tumor-specific CD8^+^ T cells exit the tumor, thereby limiting tumor control. CXCR4 inhibition and loss of lymphatic-specific CXCL12 boosts T cell retention and enhances response to therapeutic immune checkpoint blockade. Strategies that limit T cell egress, therefore, provide a new tool to boost immunotherapy response.

**One-Sentence Summary:** Lymphatic vessel-mediated, antigen-dependent CD8^+^ T cell egress limits T cell accumulation in melanomas and impairs anti-tumor immunity.

---

Lymphatic vessels are integral and active participants in the generation, maintenance, and resolution of adaptive immunity ^1^. Blind-ended lymphatic capillaries unidirectionally transport interstitial fluid (containing soluble peptides, antigens, and lipids) and leukocytes from peripheral tissues to lymph nodes (LN). Although necessary for *de novo* adaptive immunity in malignant and non-malignant settings ^2–4^, lymphatic vessels can be immune suppressive, present antigen for T cell dysfunction ^5, 6^, and limit CD8^+^ T cell-dependent tumor control in an IFNγ-dependent manner^7^. These data indicate that lymphatic vessels and their transport function may directly contribute to tumor immune evasion. However, our understanding for how lymphatic vessels influence CD8^+^ effector T cell function within the tumor microenvironment (TME) remains incomplete.

CD8^+^ T cell infiltration into the tumor parenchyma is associated with improved patient outcomes ^8^ and response to immune checkpoint blockade (ICB) ^8, 9^. The accumulation of CD8^+^ T cells in tumors is determined by mechanisms of recruitment, retention, and exit via lymphatic vessels. T cell recruitment to inflamed tissues, including tumors, is antigen-independent, resulting in the presence of both tumor-specific and irrelevant bystanders in TMEs ^10, 11^. While local antigen encounter in virally-infected tissue helps to focus the tissue resident memory repertoire ^12^, how chronic antigen and a dysfunctional tissue environment influences T cell retention in tumors remains unknown. Interestingly, CD4^+^ and CD8^+^ T cells access afferent lymph at steady state and during inflammation ^13–15^ and lymphocytes exit the TME in preclinical models ^16–18^, raising the possibility that the rate or selectivity of CD8^+^ T cell exit from TMEs might also shape the functional intratumoral repertoire and thereby response to therapy.

In this study, we investigated the hypothesis that lymphatic vessel-mediated effector CD8^+^ T cell egress limits the accumulation of a broad repertoire of functional CD8^+^ T cells that improve tumor control in combination with ICB. We demonstrate that tumor-associated lymphatic vessels direct CD8^+^ T cell sequestration at the tumor periphery, thereby increasing the probability of eventual exit in a CXCL12-CXCR4 and antigen-dependent manner. These data therefore demonstrate that the lymphatic vasculature helps tune the diversity and functional state of the intratumoral CD8^+^ T cell repertoire and these mechanisms provide new strategies for combination immunotherapy.

## Results

### Dermal lymphatic vessels limit CD8^+^ T cell accumulation in melanoma

CD8^+^ T cell exclusion from the tumor parenchyma associates with tumor immune evasion and poor response to immunotherapy ^9^. Despite being immunogenic and highly inflamed, the implantable YUMMER1.7 murine melanoma also exhibits an exclusion phenotype that might contribute to outgrowth in the immune competent setting. In these tumors, CD8^+^ T cells are sequestered at the tumor periphery (**Fig. 1A** and **B**) and close to peritumoral lymphatic vessels (**Fig. 1C**) ^7, 19^. Though CD8^+^ T cells access dermal lymphatic vessels and recirculate out of inflamed skin ^15^ and cutaneous tumors ^16, 20^, the mechanisms that govern their exit from tumors, and the functional impact of exit on tumor control remains completely unknown. Interestingly, we previously demonstrated that response to adoptive T cell therapy (ATT) was enhanced in mice lacking dermal lymphatic vessels compared to wild type (WT) littermate controls ^2^, leading us to hypothesize that functional lymphatic vessels facilitate T cell exit, thus limiting tumor-specific CD8^+^ T cell accumulation and anti-tumor immune control.

**Fig. 1.**
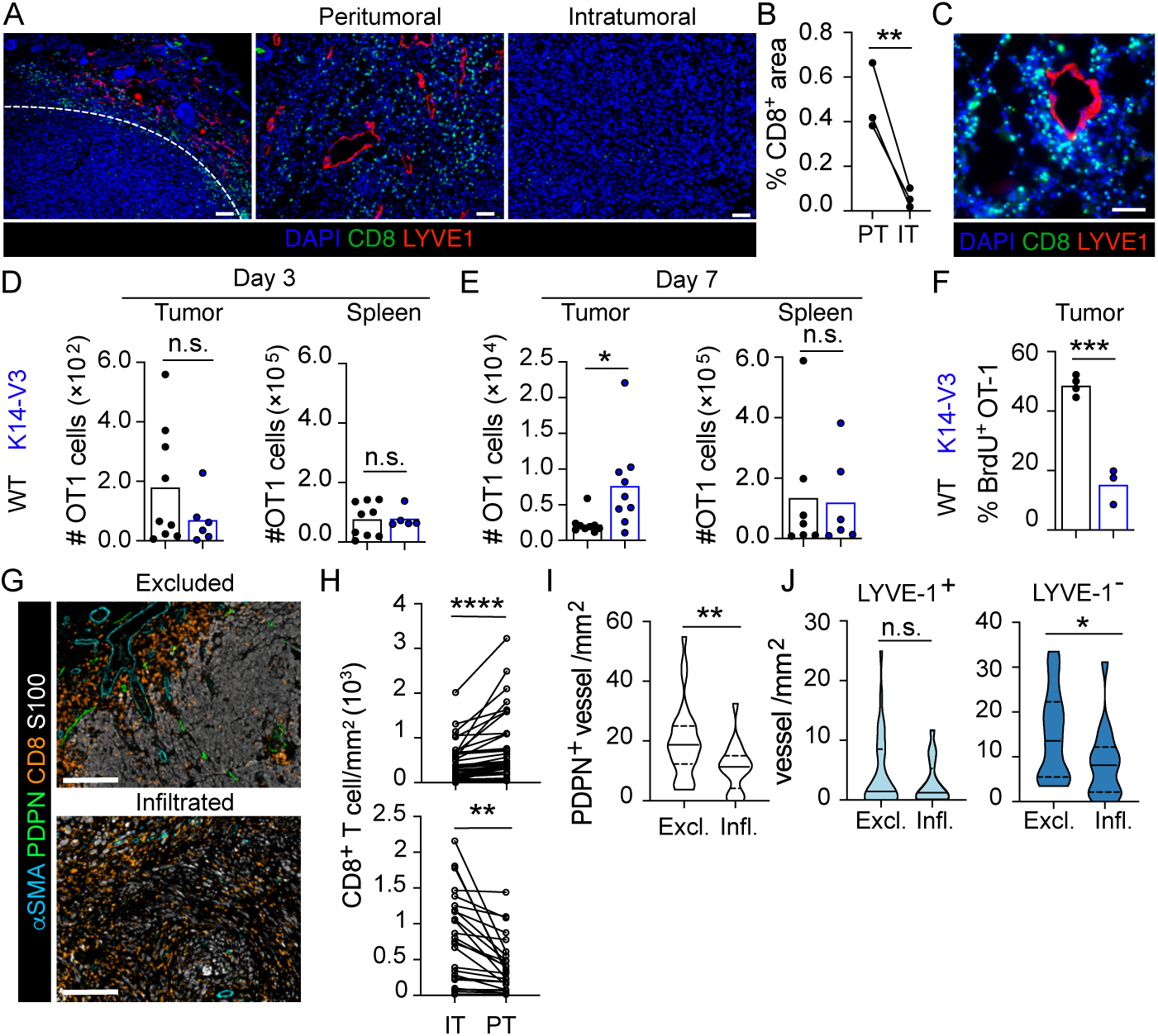
Dermal lymphatic vessels limit CD8^+^ T cell accumulation in melanoma. (**A**) Representative immunofluorescence images of peritumoral and intratumoral CD8^+^ T cells (green) and LYVE-1^+^ (red) lymphatic vessels from YUMMER1.7 tumors. Nuclei stained with DAPI. Left image: scale bar = 100 μm middle and right images: scale bar = 50 μm. (**B**) Average frequency of CD8^+^ pixels per peritumoral (PT) or intratumoral (IT) of YUMMER1.7 tumor. Each dot represents one mouse (n=3). (**C**) Representative image of CD8^+^ T cells (green) located around a LYVE-1^+^ lymphatic vessel (red). Scale bar = 50 μm. (**D-E**) 5×10^5^ congenically labeled ex vivo activated OT1 T cells were intravenously transferred into B16.F10^OVA^- bearing WT or K14-VEGFR3-Ig (K14-V3) mice on day 7. Graphs indicate numbers of Thy^1.1/1.1^ OT1 T cells in B16.F10^OVA^ tumors and spleens at day 3 (D) and day 7 (E) of WT and K14-V3 mice. n=18 WT and 15 K14-V3 across 2-3 independent experiments. (**F**) Percent BrdU^+^ of all Thy^1.1/1.1^ OT1 T cells in B16.F10^OVA^ tumors on day 7 (n=4). (**G**) Representative multiplex IHC images of T cell excluded (top image) and infiltrated (bottom image) regions in human cutaneous melanomas. αSMA^+^ blood vessels (cyan), podoplanin^+^ lymphatic vessels (PDPN, green), CD8^+^ T cells (orange), S100^+^ melanoma (white). Scale bar = 400 μm. (**H**) Number of CD8^+^ T cells per mm^2^ in the intratumoral (IT) vs peritumoral (PT) regions from (G). (**I**) Number of PDPN^+^ vessels per mm^2^ in excluded (Excl.) vs infiltrated (Infl.) regions. (**J**). Number of LYVE-1^+^ (left) or LYVE-1^-^ (right) PDPN^+^ lymphatic vessels per mm^2^ in excluded vs infiltrated regions. For G-J, across 28 patient samples, 47 ROIs were scored as “excluded” and 25 ROIs scored as “infiltrated; each symbol represents one ROI. For B and D-F, each symbol represents one mouse. Two-sided, unpaired (B-F and I-J) and paired student’s t tests (H). *p<0.05, **p<0.01, ***p<0.001, ****p<0.0001, n.s.=not significant.

To begin to understand how lymphatic vessels directly contribute to antigen-specific CD8^+^ T cell accumulation, we tracked the recruitment, accumulation, and proliferation of transferred CD8^+^ T cells *in vivo.* T cell receptor-transgenic (TCR-Tg) OT1 CD8^+^ T cells, specific for the immunodominant MHCI-restricted epitope in ovalbumin (OVA), equally infiltrated both lymphatic-deficient (K14-VEGFR3-Ig ^21^) and wildtype (WT) B16.F10^OVA^ tumors and established similar circulating frequencies three days post transfer (**Fig. 1D**). Interestingly, however, by day 7 post transfer, OT1 CD8^+^ T cells accumulated more in lymphatic-deficient melanomas relative to WT, while circulating numbers remained equal (**Fig. 1E**). This boost in OT1 CD8^+^ T cells was not explained by proliferation; in fact, we observed significantly reduced OT1 CD8^+^ T cell proliferation in lymphatic vessel-deficient TMEs (**Fig. 1F**). These data indicated that lymphatic transport limits the efficacy of ATT by decreasing retention of therapeutic T cells, despite their high affinity antigen specificity. Interestingly, CD8^+^ T cell excluded regions of human cutaneous primary melanomas (**Fig. 1G** and **H**) exhibited increased lymphatic vascular density (PDPN^+^) relative to peritumoral regions with adjacent robust tumor infiltration (**Fig. 1I**). Notably, the change in lymphatic vessel density was driven by the LYVE-1^-^PDPN^+^ subset (**Fig. 1J**). LYVE-1 loss on lymphatic endothelial cells (LEC) is driven by inflammation ^22^, raising the interesting possibility that both the presence of dermal lymphatic vessels, and their functional state, may contribute to CD8^+^ T cell accumulation and tumor control.

### Lymphatic-secreted CXCL12 sequesters CD8^+^ T cells in the tumor periphery

These data informed two complementary hypotheses, first that the inflamed state of the tumor-associated lymphatic vasculature might be poised to attract CD8^+^ T cells, and second, that CD8^+^ T cell proximity to peripheral lymphatic vessels could lead to enhanced rates of exit ^16, 20^. Tumor-associated LECs isolated from autochthonous Tyr::CreER;*Braf^V600E^;Pten*^fl/fl^ murine melanomas (BPC; **Fig. S1A** and **B**) ^23^, exhibited tumor-induced transcriptional reprogramming (**Fig. S1C-E** and **Table S1**). Tumor-associated LECs significantly upregulated genes associated with extracellular matrix remodeling and inflammation (**Fig. 2A**), including a shift in chemokine expression that could have a significant impact on leukocyte position. Interestingly, expression of *Ccl21a* and the sphingosine kinases (*Sphk*) required for S1P production were unchanged (*Ccl21a, Sphk2*) or decreased (*Sphk1*) in tumor-associated LECs compared to naïve skin (**Fig. S1F-H**). Conversely, *Cxcl12*, a chemokine implicated in dendritic cell (DC) and B cell egress ^24, 25^ and T cell exclusion ^26^, was upregulated in tumor-associated LECs relative to naive (**Fig. 2B**). *Cxcl12* was specifically expressed at the periphery of YUMMER1.7 tumors (**Fig. S2A**) in regions of high CD8^+^ T cell density (**Fig. 1A and B**). *Cxcl12*-reporter mice revealed that CD45^-^ stromal cells were major producers of *Cxcl12* in YUMMER1.7 tumors (**Fig. S2B**). Furthermore, 16.6% (±6.9) of LECs (**Fig. S2C** and **D**) and 29.4% (±8.8) of blood endothelial cells (BECs) expressed *Cxcl12*, while only a minor percentage of all epithelial cells, fibroblasts, and immune cells showed significant expression (**Fig. S2D** and **E**). A similar distribution of expression was also observed in the non-immunogenic YUMMER1.7 parental line, YUMM1.7 (**Fig. S2F-H**). Interestingly, the frequency of *Cxcl12*^+^ BECs and fibroblasts was lower in YUMMER1.7 tumors compared to normal skin (**Fig. S2I** and **J**), while *Cxcl12*^+^ LECs were more abundant in these same tumors (**Fig. 2C**), and largely LYVE-1 negative (**Fig. 2D**), which is consistent with the lymphatic vessel phenotype enriched in excluded human melanomas (**Fig. 1J).** Immunofluorescence further confirmed the abundance of CXCL12 in the tumor periphery and its association with a subset of tumor-associated lymphatic vessels (*Prox1*-tdTomato^+^, **Fig. 2E**).

**Fig. 2.**
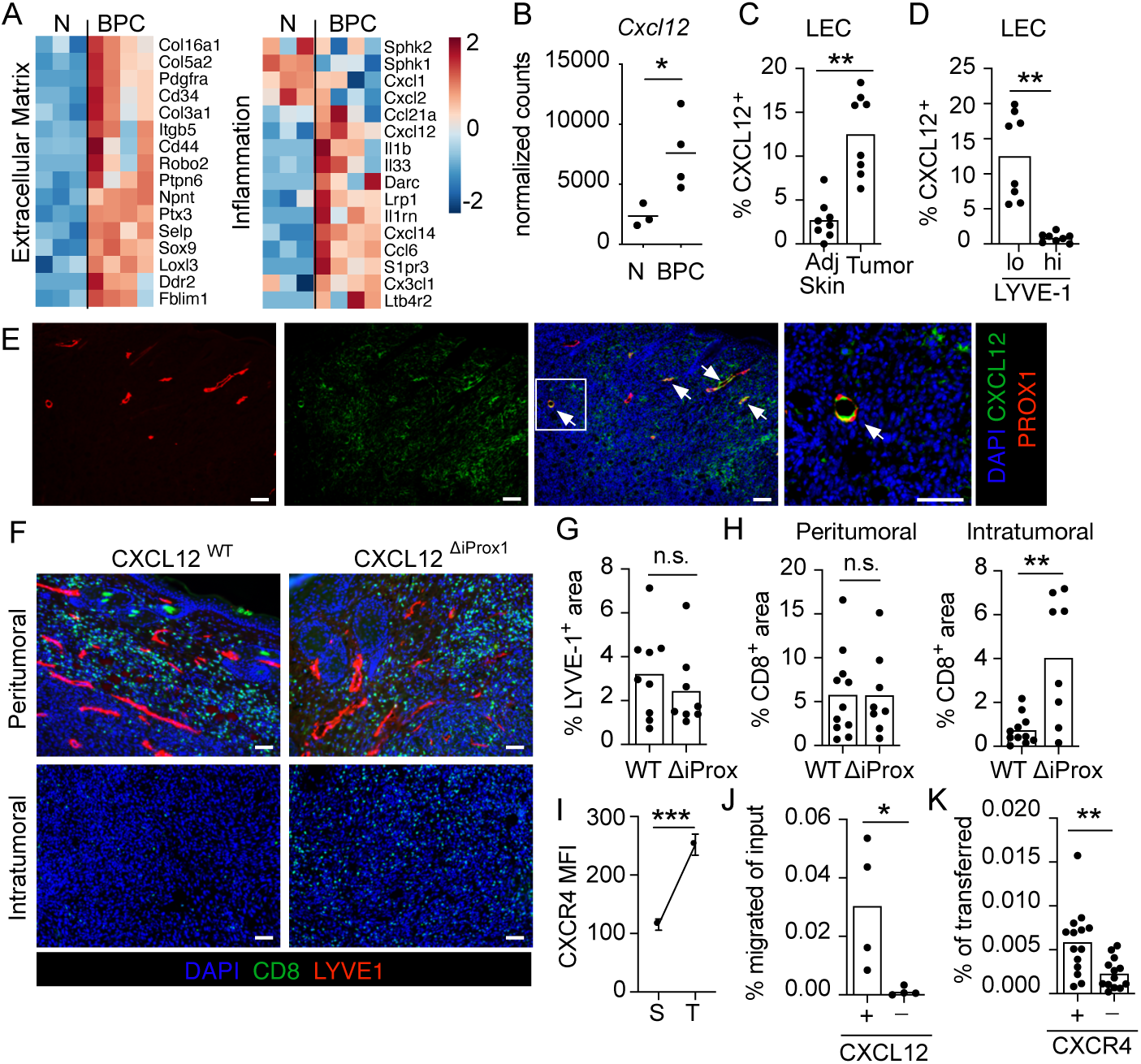
Lymphatic endothelial cell-secreted CXCL12 sequesters CD8^+^ T cells in the tumor periphery. (**A**) Heatmap of select genes differentially expressed by sorted CD45^-^CD31^+^gp38^+^ lymphatic endothelial cells (LEC) from naïve skin (N; n= 3 mice) or BPC murine melanomas (BPC; n= 4 mice). (**B**) Normalized CXCL12 transcript counts from naïve or BPC tumor-associated LECs. (**C**) Frequency of CXCL12-dsRed^+^ cells of CD45^-^CD31^+^gp38^+^ LECs from normal adjacent skin or YUMMER.17 tumors in CXCL12-dsRed reporter mice. (**D**) Frequency of CXCL12-dsRed^+^ cells of gp38^+^LYVE-1^lo^ or gp38^+^LYVE-1^hi^ LECs. For C-D, n=8. (**E**) Representative images of CXCL12 staining (green) in YUMMER1.7 tumors implanted in *Prox1*-TdTomato (red) reporter mice. Nuclei, DAPI (blue). Arrows = CXCL12^+^ lymphatic vessels; scale bar = 100 μm. Inset, far right; scale bar = 50 μm. (**F**) Representative peritumoral and intratumoral images of CD8^+^ T cells (green) and LYVE-1^+^ (red) lymphatic vessels in YUMMER1.7 tumors (day 14) of CXCL12^WT^ or CXCL12^ΔiProx1^ mice. Nuclei, DAPI (blue). Scale bars = 50 μm. (**G)** Frequency of LYVE-1^+^ pixels per image in the peritumoral region of YUMMER1.7 tumors in CXCL12^WT^ (n=9) or CXCL12^ΔiProx1^ (n=8) mice. (**H**) Frequency of CD8^+^ pixels per image in peritumoral and intratumoral regions of YUMMER1.7 tumors in CXCL12^WT^ (n=11) or CXCL12^ΔiProx1^ (n=8) mice; results of 2 independent experiments. (**I**) CXCR4 mean fluorescence intensity (MFI) in CD44^+^CD8^+^ T cells in the spleen (S) or YUMMER1.7 tumor (T). Paired student’s t-test; n= 3; error bar = standard deviation. (**J**). Frequency of CD44^+^CD8^+^ T cells migrated toward serum free media (-) or 100 ng/ml CXCL12 (+). n=4 mice. (**K**) Frequency of wild type (+) (n=14) or CXCR4-deficient (-) (n=13) effector CD45.2^+^CD8^+^ T cells in tumor dLNs 20hrs post intratumoral transfer. For K, 3 independent experiments were performed. For all experiments, each symbol represents one mouse. Two-sided, unpaired (B, G-H, and J-K) and paired (C-D and I) student’s t-tests. *p<0.05, **p<0.01, ***p<0.0001, n.s.=not significant.

To uncover the contribution of LEC-derived CXCL12 to CD8^+^ T cell accumulation and position within the melanoma TME, we crossed the inducible *Prox1:Cre^ERT2^* with *Cxcl12*^fl/fl^ mice to ablate *Cxcl12* from LECs (*Cxcl12*^ΔiProx1^) (**Fig. S2K**). LEC-specific *Cxcl12* deletion did not impact peritumoral lymphatic vessel density (**Fig. 2F** and **G**), however, it appeared to dramatically enhance CD8^+^ T cell infiltration into the tumor parenchyma (**Fig. 2F** and **H**), indicating that independent of changes in vessel number, chemokines derived from activated peritumoral lymphatic vessels contribute to T cell position within the TME. Importantly, intratumoral CD8^+^ T cells expressed higher levels of surface CXCR4 than their circulating counterparts (**Fig 2I** and **Fig. S3A)** and effector CD8^+^ T cells were sensitive to CXCL12 gradients *ex vivo* (**Fig. 2J**). Based on these observations, we considered the possibility that lymphatic-derived inflammatory chemokines, and in particular CXCL12, were directing CD8^+^ T cells to the tumor periphery where they would, in turn, be more likely to exit the TME ^16, 20^. Indeed, while intratumorally transferred CD8^+^ T cells accessed the tumor draining LN (dLN), CXCR4 knockout (CXCR4^ΔUBC^; **Fig. S3B** and **C**) effector CD8^+^ T cells were significantly impaired in their ability to egress from the TME (**Fig. 2K**). These data together indicate that lymphatic vessel-derived CXCL12 in part regulates the probability of CXCR4^+^ T cell exit from tumors.

### Functional tumor-specific CD8^+^ T cells egress from tumors via lymphatic vessels

In order to understand the functional consequence of CD8^+^ T cell egress on the accumulation and diversity of the intratumoral repertoire we leveraged a photoconvertible Kaede-Tg mouse model to track endogenous leukocyte migration *in vivo* ^17^. We found that CD8^+^ T cells represented 25-30% of egressing leukocytes across several implantable models including melanoma e.g. YUMM1.7 (22.35% ±7.9), YUMMER1.7 (24.98% ±6.4) and B16.F10 (28.43% ±5.2), colorectal (MC38) and fibrosarcoma (MCA.205) (**Fig. 3A** and **Fig. S3D** and **E**), and consistent with a role for CXCR4 in T cell egress, pertussis toxin, but not FTY720, inhibited CD8^+^ T cell egress (**Fig. S3F-H)**. Egressing CD8^+^ T cells were preferentially CD44^+^CD62L^+^ central memory, while CD44^+^CD62L^-^ effector/effector memory cells were largely retained (**Fig. S4A-B**), consistent with the known circulating properties of these T cell subsets.

**Fig. 3.**
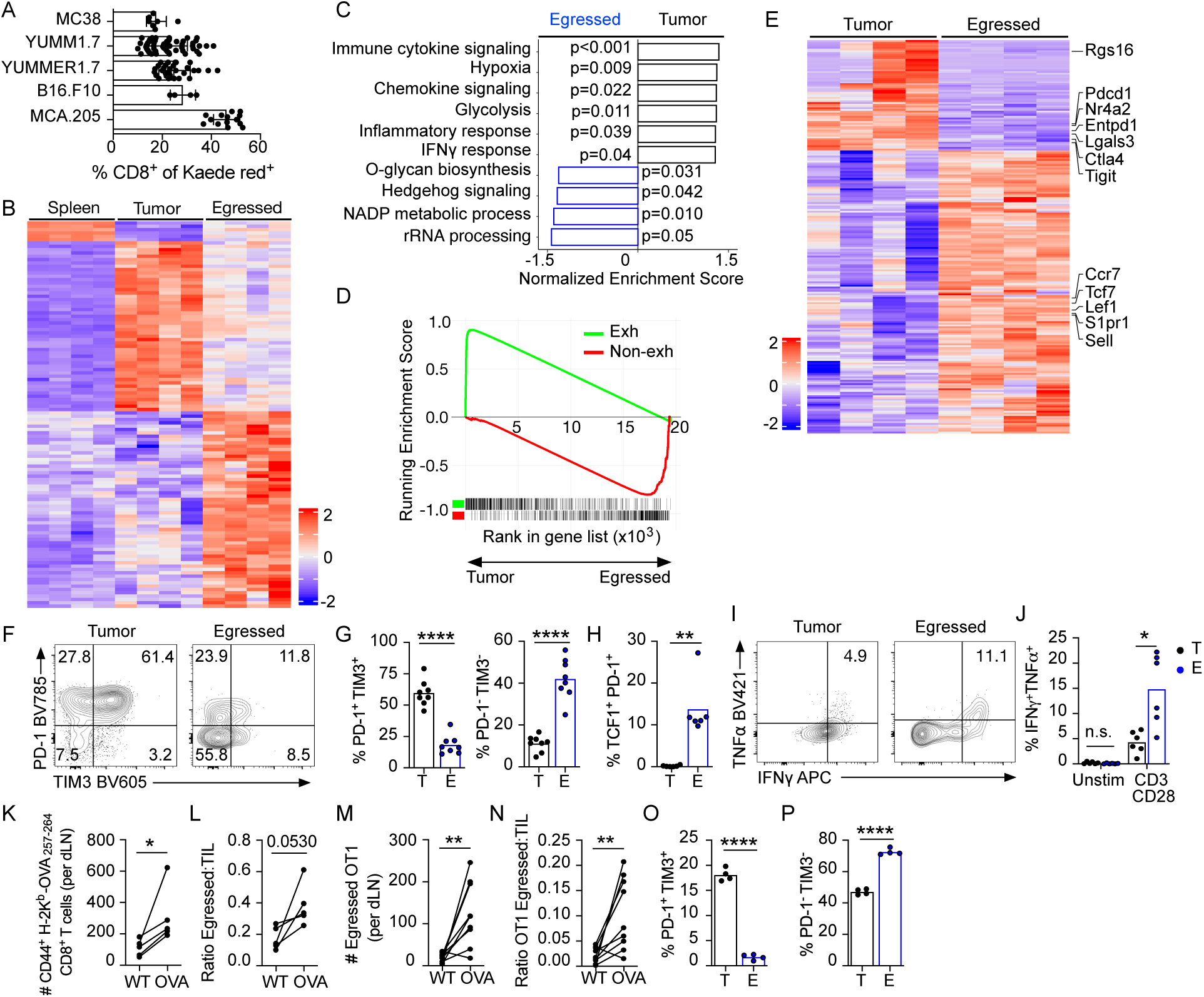
Functional tumor-specific effector CD8^+^ T cells egress from tumor via lymphatic vessels. (**A**) CD8^+^ T cell frequency of Kaede red^+^ cells in the dLNs of various tumors 24hrs post-photoconversion of Kaede-Tg mice. MC38: n=7; YUMM1.7: n=54; YUMMER1.7: n=30; B16.F10: n=4; MCA.205: n=14; except for B16.F10 tumors, 2 or more independent experiments were performed; error bars = standard deviation. (**B**). Heatmap of differentially expressed genes in circulating (spleen), tumor retained (Kaede red^+^), and egressed (Kaede red^+^) CD44^+^CD8^+^ T cells identified by RNA-Seq. n=4 mice (**C**) Gene set enrichment analysis of pathways enriched in tumor vs egressed CD44^+^CD8^+^ T cells. (**D**) Enrichment scores for gene signatures associated with T cell dysfunction (GSE41867) in tumor versus egressed CD44^+^CD8^+^ T cells; up in exhausted (Exh, green); down in exhausted (Non-exh, red). (**E**) Heatmap of differentially expressed leading edge genes from GSE41867 in tumor versus egressed CD44^+^CD8^+^ T cells. (**F**) Representative PD-1 and TIM3 flow plots on intratumoral and egressed CD44^+^CD8^+^ T cells from YUMMER1.7 tumors. (**G-H**) Frequencies of PD-1^+^TIM3^+^, PD-1^-^TIM3^-^ (G; n=8) and TCF1^+^PD-1^+^ (H; n=6) cells among tumor (T) and egressed (E) CD44^+^CD8^+^ T cells of YUMMER1.7. (**I-J**) Representative flow plots (I) and frequency (J) of *ex vivo* IFN and TNF production by CD44^+^CD8^+^ T cells tumor retains (T) or egressed (E) YUMMER1.7 tumors (day 21). n=6 across 2 independent experiments. (**K-L**) Numbers (K) and egress rate (ratio of number of egressed to number in tumor; (L)) of H-2K^b^ OVA_257-264_ CD44^+^CD8^+^ T cells to dLNs of MCA.205^WT^ (WT) or MCA.205^OVA^ (OVA) tumors. n=5. (**M-N**) 4×10^6^ *in vivo* activated OT1-Kaede-Tg were transferred into mice bearing B16.F10^WT^ and B16.F10^OVA^ on opposite flanks (day 7). Three days post transfer, both tumors were photoconverted. Graphs indicate numbers (M) and egress rate (N) of Thy^1.1/1.1^CD44^+^ OT1 T cells to dLNs of B16.F10^WT^ or B16.F10^OVA^ tumors 24hrs post photoconversion. n=9 across 2 independent experiments. (**O-P**) Frequencies of PD-1^+^TIM3^+^ (O) and PD-1^-^ TIM3^-^ (P) cells of tumors (T) or egressed (E) Thy^1.1./1.1^CD44^+^ OT1 T cells of B16.F10^OVA^ tumors. n=4. For all experiments, each symbol represents one mouse. Two-sided, unpaired (G, H, J) and paired (K-P) student’s t-tests. *p<0.05, **p<0.01, ****p<0.0001.

To define the transcriptional programs associated with egress, we sorted Kaede red^+^ CD44^+^CD8^+^ T cells from dLNs (egressed) and YUMM1.7 tumors (retained) as well as circulating CD44^+^CD8^+^ T cells from the spleen and performed bulk RNAseq. Egressed effector CD8^+^ T cells were transcriptionally distinct from both their retained and circulating counterparts (**Fig. 3B** and **Table S2**). While tumor-retained effector CD8^+^ T cells were enriched for transcriptional programs associated with hypoxia, chemokine and cytokine signaling, and IFNγ responses, egressed CD8^+^ T cells were enriched for O-glycan biosynthesis, ribosomal RNA processing, and hedgehog signaling (**Fig. 3C**). Interestingly, genes associated with CD8^+^ T cell dysfunction ^27^ were notably higher in retained CD8^+^ T cells relative to recent emigrants. We therefore queried the specific association of transcripts upregulated or downregulated in dysfunctional CD8^+^ T cells ^28^ and observed a striking segregation of the transcriptional states of these two populations (**Fig. 3D**). Retained, intratumoral CD8^+^ T cells expressed transcripts upregulated in dysfunctional CD8^+^ T cells (*Pdcd1*, *Nr4a2*, *Ctla4*, and *Tigit*) and conversely, egressed CD8^+^ T cells were enriched for transcripts (*Tcf7*, *Sell*, *Lef1*, *Ccr7,* and *S1pr1)* downregulated during dysfunction and associated with migration and a stem-like phenotype (**Fig. 3D-E** and **Table S3**).

The transcriptional data indicated phenotype selection as a function of CD8^+^ T cell egress and retention. In concordance, intratumoral effector CD8^+^ T cells in YUMMER1.7 (and YUMM1.7) tumors were predominantly PD-1^+^TIM3^+^ and LAG3^+^ while egressed CD8^+^ T cells lacked expression of all of these molecules (**Fig. 3F-G** and **Fig. S4C-H**). Moreover, egressed CD8^+^ T cells were predominantly TCF1^+^ (**Fig. 3H** and **Fig. S4I** and **J**) consistent with a circulating central memory population; while intratumoral CD8^+^ T cells were largely TCF1^-^. Interestingly, however, a subset of egressed CD8^+^ T cells expressed both TCF1 and PD-1 (**Fig. 3H**), consistent with a stem-like CD8^+^ progenitor population ^29, 30^ thought to drive therapeutic response to ICB ^31^ and found proximal to lymphatic vessels in renal cancer ^19^. These data raised the possibility that a functional pool of CD8^+^ T cells was poorly retained in tumors, limiting effective tumor control. Concordantly, egressed CD8^+^ T cells from YUMMER1.7 (**Fig. 3I** and **J**) and YUMM1.7 (**Fig. S4K** and **L**) tumors robustly produced effector cytokines upon *ex vivo* restimulation while retained CD8^+^ T cells did not.

To determine if a subset of these functional, egressing CD8^+^ T cells could be tumor-specific, we utilized an immunogenic tumor line, MCA.205, engineered to express OVA (MCA.205^OVA^) and its isogenic wildtype counterpart (MCA.205^WT^). Each tumor line was co-implanted on opposite flanks of Kaede-Tg mice and endogenous H-2K^b^-OVA_257-264_ T cell egress was measured from each tumor to their respective dLNs on day 9. As expected, OVA-specific CD8^+^ T cells egressed from MCA.205^WT^ tumors lacking OVA expression. We also observed, however, significant OVA-specific CD8^+^ T cell egress from MCA.205^OVA^ tumors (**Fig. 3K-L** and **Fig. S5A**). We similarly evaluated tumor-specific CD8^+^ T cell egress by adoptively transferring Kaede-expressing exogenous OT1 TCR-Tg CD8^+^ T cells into mice bearing B16.F10^WT^ and B16.F10^OVA^ tumors. Even when expressing a fixed, high affinity TCR, we still observed significant CD8^+^ egress in the presence of cognate antigen (B16.F10^OVA^; **Fig. 3M-N** and **Fig. S5B**). Unbiased TCRβ sequencing of endogenous intratumoral and egressed CD3ε^+^ T cells revealed clonal overlap (shared clones; **Fig. S6A-C** and **Table S4**). The clonal frequency of these shared clones was between 0.01-1.0% (medium to large) indicating clonal expansion within tumors (**Fig. S6D-F**) and consistent with the hypothesis that a subset of tumor-specific T cells exit the TME. Importantly, egressed OT1 CD8^+^ T cells phenocopied the polyclonal population (**Fig 3F-G**), and were more likely to be PD-1^-^TIM3^-^ while tumor-retained OT1 CD8^+^ T cells were more likely to be PD-1^+^TIM3^+^ (**Fig. 3O** and **P** and **Fig. S5C**), suggesting that strategies to inhibit egress might improve functional, antigen-specific CD8^+^ T cell retention and thereby response to immunotherapy.

### Antigen encounter in the tumor microenvironment links T cell function and migratory potential

Local antigen encounter in tumors regulates exhaustion ^32, 33^ and retention ^12^. Our *in vivo* data indicated that lymphatic-derived CXCL12 restricts CD8^+^ T cells to the tumor periphery and that CXCR4 expression in CD8^+^ T cells enhanced their migration to dLN. Given the observed differences in egress between dysfunctional and functional CD8^+^ T cell states in our murine models (**Fig. 3**), we hypothesized that antigen encounter might simultaneously tune surface expression of CXCR4 and intrinsic T cell function. Consistent with this hypothesis, CXCR4 was highly expressed on PD-1^-^TIM3^-^ CD8^+^ T cells but low on dysfunctional PD-1^+^TIM3^+^ within YUMMER1.7 tumors (**Fig. 4A** and **B**). Conversely, known regulators of T cell retention and tissue residency, such as CXCR6, were significantly enriched on PD-1^+^TIM3^+^ retained CD8^+^ T cells (**Fig. 4C**). Interestingly, we also observed a positive correlation between surface ACKR3 (CXCR7), a decoy receptor for CXCL12, and the dysfunctional PD-1^+^TIM3^+^ subset (**Fig. 4D**), providing a second mechanism through which antigen encounter tunes CXCL12 sensitivity *in vivo*. To validate these relationships between CD8^+^ T cell state and chemokine receptor expression in human melanoma, we analyzed an existing CD8^+^ T cell single cell RNA-Seq dataset from untreated and ICB-treated patients ^34^ and defined five cell states consistent with the literature (**Fig. S7**) ^27^. *CCR7* and *S1PR1* expression was highest in memory/early activated CD8^+^ T cells (*CCR7*^+^*SELL*^+^*TCF7*^+^*IL7R*^+^) and lowest in pre-dysfunctional (*TCF7*^+^*NR4A3*^+^*CTLA4*^+^) and exhausted (*TCF7*^-^*PDCD1*^+^*CTLA4*^+^*LAG3*^+^) T cells (**Fig. 4E**). Conversely, residency genes *RGS1* and *CXCR6* ^35, 36^ were highly expressed by exhausted and late exhausted (*TCF7*^-^ *PDCD1*^hi^*CTLA4*^hi^*LAG3*^hi^) CD8^+^ T cells populations (**Fig. 4E**). *CXCR4*, which is constitutively expressed by CD8^+^ T cells, was lowest in cytotoxic (*CCR7*^-^*IL7R*^-^*PRF1*^+^) and exhausted CD8^+^ T cells (**Fig. 4E**), states that were least likely to egress in our murine models (**Fig. 3F** and **G**), and highly expressed by early-activated /memory and pre-dysfunctional CD8^+^ T cells, states most likely to egress (**Fig. 3G** and **H**).

**Fig. 4.**
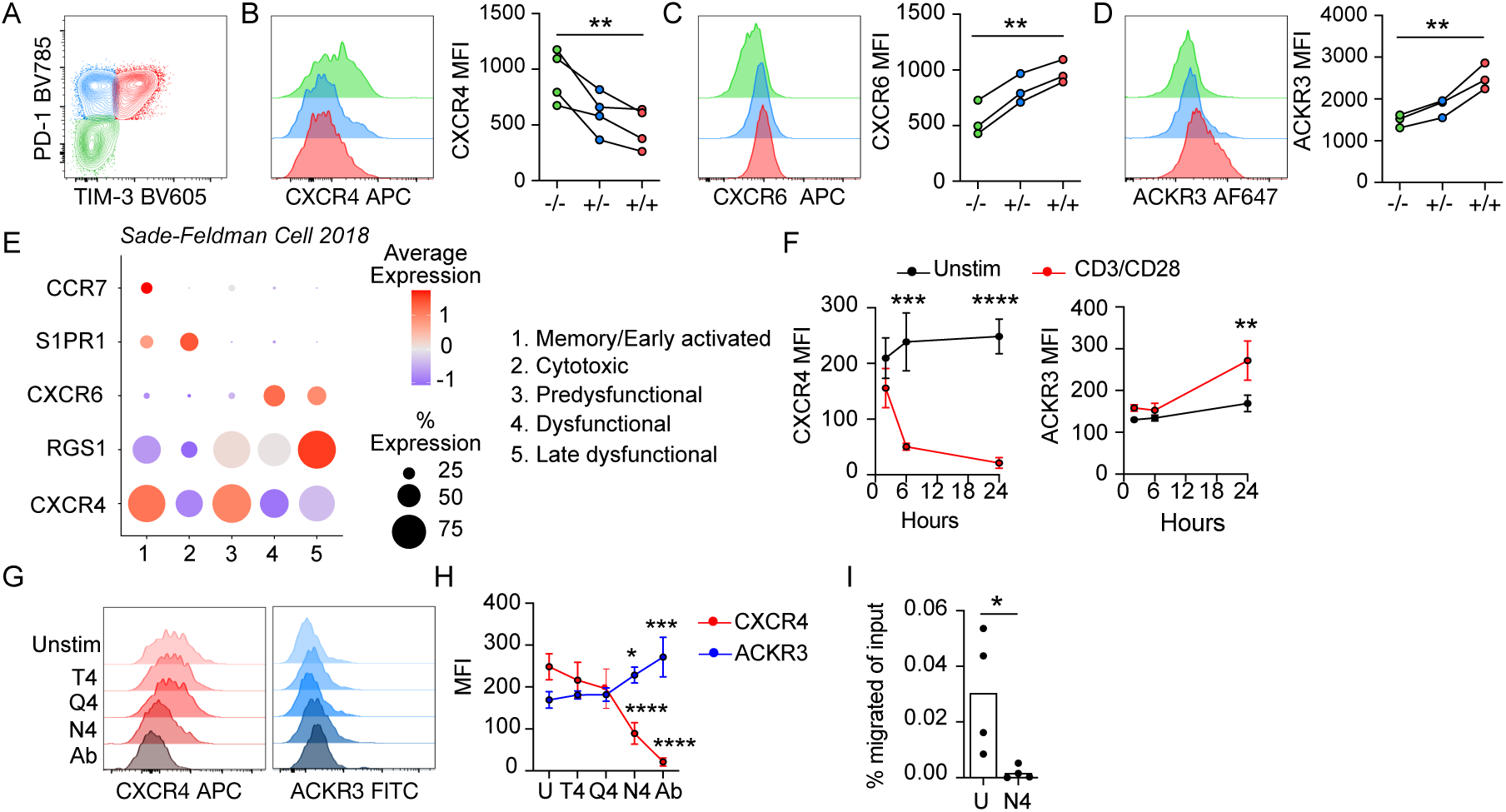
Antigen encounter in the tumor microenvironment links function and migratory potential. (**A**) Representative flow plot identifying the dysfunctional states of intratumoral CD44^+^CD8^+^ T cells used in figures B-D: PD-1^-^TIM3^-^ (green; -/-); PD-1^+^TIM3^-^ (blue; +/-); PD-1^+^TIM3^+^ (red; +/+). (**B-D**) Surface expression of CXCR4 (B), CXCR6 (C), and ACKR3 (D) on intratumoral PD-1 vs TIM3 CD44^+^CD8^+^ T cells from YUMMER1.7 tumors. For B-D, n=3 or 4 mice; each symbol represents one mouse. (**E**) Bubble plot demonstrating the expression of select genes regulating T cell migration across T functional states within human melanomas (GSE120575). Size of the dot represents the percentage of cells expressing the gene of interest in each functional group; color of dot represents the average level of expression of the transcript. (**F**) Time course of CXCR4 (left) and ACKR3 (right) surface expression on OVA-specific CD44^+^CD8^+^ T cells following *ex vivo* stimulation in the absence (unstim) or presence of CD3/CD28 antibodies. (**G-H**) Representative histograms (G) and surface expression (H) of CXCR4 and ACKR3 on OVA-specific CD44^+^CD8^+^ T cells following 24hr *ex vivo* stimulation in the absence (U) or presence of CD3/CD28 antibodies (Ab) or peptides: SIINFEKL (N4), SIITFEKL (T4), or SIIQFEKL (Q4). For F-H, n=4 mice; representative graph from 3 independent experiments. (**I**) Frequency of unstimulated (U) or SIINFEKL-stimulated (N4) OVA-specific CD44^+^CD8^+^ T cells that migrated to CXCL12. n=4 mice; representative graph of 2 independent experiments. MFI=mean fluorescence intensity. One-way ANOVA adjusted for multiple comparisons (B-D, F, and H); two-sided, unpaired student’s t-test (I). *p<0.05, **p<0.01, ***p<0.001, ****p<0.0001.

We therefore evaluated the relationship between antigen encounter by infiltrating effector CD8^+^ T cells and their potential to migrate along a CXCL12 gradient. *Ex vivo* TCR stimulation of effector CD44^+^CD8^+^ T cells with CD3ε/CD28 antibodies significantly and rapidly reduced CXCR4 surface levels, while ACKR3 surface expression increased, reaching peak expression 24 hours post-stimulation (**Fig. 4F**). Given the overlap in the TCR repertoire (**Fig. S6**) and the impact of suboptimal antigen encounters on T cell phenotype in tumors ^37^, we explored the impact of antigen affinity on chemokine receptor surface expression. Endogenous OVA-specific effector CD44^+^CD8^+^ T cells were stimulated with the immunodominant epitope of OVA, SII<u>N</u>FEKL (N4; potency=1), or peptides with single amino acid substitutions that elicit weaker TCR signaling: SII<u>Q</u>FEKL (Q4; potency 1/18 of N4) or SII<u>T</u>FEKL (T4; potency 1/71 of N4) ^38^. Weak TCR signaling (Q4 and T4) failed to reduce CXCR4 or increase ACKR3 surface expression on CD44^+^CD8^+^ T cells (**Fig. 4G** and **H**), despite activating cytokine production (**Fig. S8A**), indicating that surface CXCR4 and ACKR3, and thus CD8^+^ T cell sensitivity to CXCL12, is tuned by the strength of TCR stimulation. As tumors present an array of neoantigens and endogenous antigens of varying affinities, this tunable expression of chemokine receptors may shape the tumor-specific, intratumoral repertoire through changes in sensitivity to competing chemokine gradients. Consistent with this hypothesis, high affinity TCR stimulation was sufficient to impede effector CD8^+^ T cell migration toward CXCL12 *ex vivo* (**Fig. 4I**). These data together indicate that the migratory fate of a CD8^+^ T cell within the TME depends not only on the stochastic probability that it will encounter its antigen, but the strength of that antigen encounter itself.

### Inhibition of the CXCL12/CXCR4 axis improves ICB tumor control in immunogenic melanoma

If CXCR4/CXCL12 signaling is causally linked to the depletion of antigen-specific CD8^+^ T cells from the TME, it stands to reason that targeting this axis would boost intratumoral T cell accumulation and prime the TME for immunotherapy. Thus, we used AMD3100 to pharmacologically inhibit CXCR4 in YUMMER1.7 melanoma. Unlike FTY720, an acute regimen of AMD3100 did not lymphodeplete the blood or lymph (**Fig. S8B**), indicating that CXCR4 is dispensable for lymphocyte recirculation from lymphoid organs. Continued administration of AMD3100 (7 days), however, increased numbers of CD45^+^ leukocytes within the TME (**Fig. 5A**) including CD3 ^+^ T cells and CD44^+^CD8^+^ T cells (**Fig. 5B** and **C**). Furthermore, TCR sequencing of intratumoral CD3 ^+^ T cells revealed that although there was no difference in the overall diversity of effector T cells within the tumor (**Fig. S8C** and **Table S5**), the maximum clonal frequency was higher with AMD3100 treatment (**Fig. 5D**), which was partially driven by an increased frequency of hyperexpanded clones (**Fig. 5E**). As expected, CXCR4 inhibition had no effect on the dysfunctional phenotype acquired by retained lymphocytes *in vivo* (**Fig. 5F**), nor did CXCR4/CXCL12 signaling directly impact effector T cell function *ex vivo* (**Fig. S8D-F**).

**Fig. 5.**
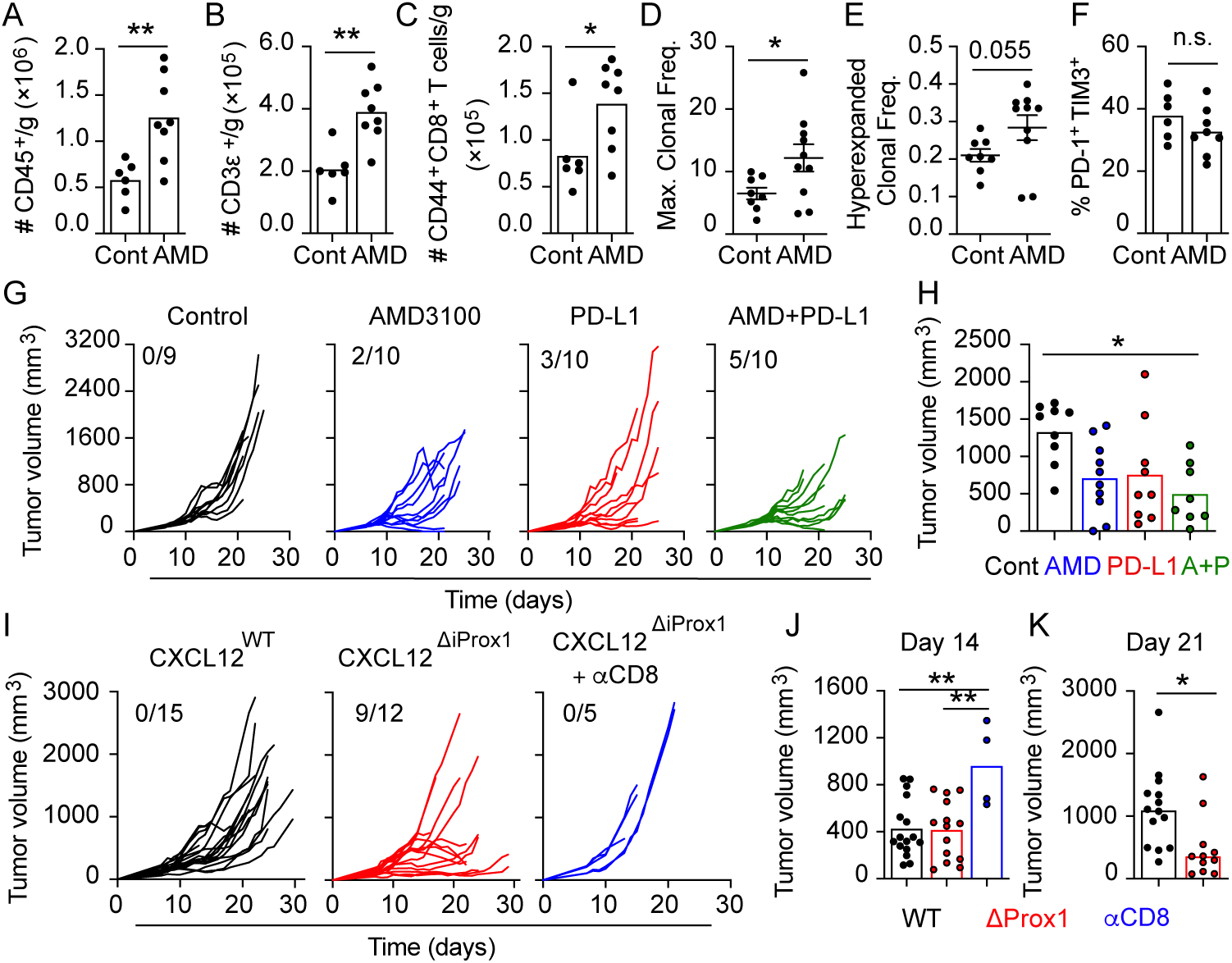
CXCL12-CXCR4 signaling blockade improves tumor control and immunotherapy efficacy. (**A-C**) Number of CD45^+^ leukocytes (A), CD3 ^+^ T cells (B), or CD44^+^CD8^+^ T cells (C) per gram of tumor after 7 days of treatment with AMD3100 (AMD; n=8)) or vehicle control (Cont; n=6) in mice bearing YUMMER1.7 tumors. (**D-E**) Maximal clonal frequency of endogenous CD3 ^+^ T cells (D) and relative levels of hyperexpanded CD3 ^+^ clones (E) in YUMMER.17 tumors 7 days post treatment with AMD3100 (n=10) or vehicle control (n=8). (**F**) Frequency of PD-1^+^TIM3^+^ CD44^+^CD8^+^ T cells in YUMMER1.7 tumors 7 days post treatment with AMD3100 (n=8) or vehicle control (n=6). (**G**) YUMMER1.7 tumor growth during treatment with vehicle control (Cont; black; n=10), AMD3100 (AMD; blue; n=10), PD-L1 (PD-L1; red; n=10), or combination AMD3100+ PD-L1 (A+P; green n=10). Each line represents one mouse; 2 independent experiments. (**H**) YUMMER1.7 tumor volumes at day 21 of treatment. (**I**) YUMMER1.7 tumor growth in CXCL12^WT^ or CXCL12^ΔiProx1^ with and without CD8 -depleting antibody (n=5) treatment. **(J-K**) YUMMER1.7 tumor volumes at day 14 (J) and day 21 (K) in CXCL12^WT^ or CXCL12^ΔiProx1^ mice. Each symbol or line represents one mouse. Two-sided, unpaired student’s t-test (A-C, F, and K), Mann-Whitney test (D-E), and One-way ANOVA (H). *p<0.05, **p<0.01, n.s.=not significant.

The changes in the endogenous repertoire driven by CXCR4 inhibition suggested that blocking CD8^+^ T cell egress would prime the TME for response to ICB by increasing the size and breadth of the antigen-specific pool available for reinvigoration. To test the therapeutic impact of CXCR4 inhibition in the context of ICB, we treated mice bearing established YUMMER1.7 tumors at a size (>200 mm^3^) when ICB is not curative as a single agent. Single agent AMD3100 led to a modest, but not statistically significant, slowing of tumor growth compared to control mice, but the combination of AMD3100 with anti-PD-L1 drove significant tumor control over either single agent alone (**Fig. 5G-H**). While AMD3100 may have many effects on diverse cell types, we importantly next asked if the specific loss of lymphatic-derived CXCL12 was sufficient to improve tumor control. Surprisingly, CXCL12^ΔiProx1^ mice exhibit spontaneous, CD8^+^ T cell-dependent YUMMER1.7 tumor control while tumors in CXCL12^WT^ mice grew progressively (**Fig. 5I-K**). This data therefore indicates that lymphatic vessel-derived CXCL12 is a mechanism of tumor immune evasion that limits the accumulation of functional CXCR4^+^CD8^+^ T cells in melanoma and may be targeted to enhance response to ICB. Moreover, our studies implicate the tumor-associated lymphatic vasculature as an active regulator of immune evasion by mediating CD8^+^ T cell egress from TMEs and is a novel target for combination immunotherapy.

## Discussion

The regulated circulation of T lymphocytes through the lymphatic and vascular systems is critical for immune surveillance and tissue homeostasis, and mechanisms that enhance or disrupt T cell homing are implicated across the spectrum of disease. Lymphatic vessels recirculate both memory and effector CD4^+^ and CD8^+^ T cells from peripheral tissues to dLNs and supply chemotactic signals that drive their directional migration ^39^. Interestingly, during both acute and chronic inflammation, the number of afferent lymph-borne T cells increases ^15^, indicating that the transport of T cells via lymph may help to manage peripheral tissue inflammation and immunopathology ^3, 40^. We and others have shown that cytotoxic CD8^+^ T cells also use afferent lymphatic vessels to exit tumors ^16, 17^, and yet the mechanisms driving lymphatic-mediated T cell exit and its functional implications for anti-tumor immunity remain poorly understood. In this study, we explored the mechanisms that determine lymphatic vessel-mediated egress and the functional consequence on intratumoral T cell repertoires and tumor control in melanoma. We demonstrate that peritumoral lymphatic vessels instruct the position and retention of CD8^+^ T cells in the TME in part through expression of CXCL12, which recruits and ultimately expels a broad repertoire of functional tumor-specific CXCR4^+^CD8^+^ T cells.

Lymphatic vessels and their transport function are critical for *de novo* immune surveillance^3^. As such, the expansion ^5, 41, 42^ or complete absence ^2, 4^ of tumor-associated lymphatic vessels is associated with so-called “hot” and “cold” tumors, respectively, in both preclinical models and human analyses. Based on these data, new vaccine strategies leverage lymphatic vessels to boost antigen delivery to LNs and prime protective adaptive immune responses ^43^. In the setting of inflamed malignant lesions, however, lymphatic vessels may also contribute to local immune suppression and limit therapeutic response. Tumor-associated lymphatic vessels can act as antigen-presenting cells to induce CD8^+^ T cell dysfunction ^5, 44^ and expand regulatory T cells ^6^, activate inhibitory ligand expression in response to cytotoxic activity ^7^, and induce immune suppression in LNs ^45^, altogether dampening local tumor control. In addition, our study now indicates that lymphatic vessels remove tumor-specific effector CD8^+^ T cells from tumors and thereby limit their accumulation and target engagement. These observations, that the impact of lymphatic vessel function on immunity might be contextual and adaptive, indicate that rather than relying on strategies to block or expand tumor-associated lymphatic vessels, efforts to decouple the immune-supporting and immune-suppressive mechanisms of lymphatic vessels may provide enhanced benefit for immunotherapy.

Here, we demonstrate that a subset of inflamed (LYVE-1^-^) tumor-associated lymphatic vessels produce CXCL12 and thereby direct CD8^+^ T cells to the tumor periphery where they are more likely egress. While we have not identified the tumor-dependent mechanisms that upregulate CXCL12 in LECs, hypoxia is sufficient to enhance CXCL12 expression *in vitro* ^46^. A LYVE-1^-^ CXCL12^+^ LEC phenotype may therefore be a function of the immunosuppressive TME, and in concordance, lymphatic vessels adopt unique activation states and chemokine repertoires as a function of their cytokine environment *in vivo* ^7, 47^. Still, there is limited data linking these adaptive responses in peripheral lymphatic vessels to the circulating behavior of leukocytes during active inflammation. Our data indicates that CXCL12 promotes effector and memory CD8^+^ T cell egress from melanoma. Though CXCR4 is dispensable for naïve CD8^+^ T cell egress from inflamed skin the mechanisms employed by naïve and antigen-experienced CD8^+^ T cells may be distinct ^48^. Furthermore, though CCL21/CCR7 is necessary for T cell egress from acutely inflamed skin and lung, it is dispensable for egress during chronic inflammation ^15^ and tumors ^20^, supporting the hypothesis that the dominant mechanisms of egress may be context dependent. More work is needed, however, to fully elucidate the tissue-specific and context-dependent mechanisms that drive T cell emigration and it is likely that multiple chemokines, lymphatic-derived and otherwise, collaborate to regulate this process.

Importantly, while it was possible that T cells egressing via lymphatic vessels would represent irrelevant bystanders ^10^, *in vivo* tracking of CD8^+^ T cells of known specificity revealed exit of tumor-specific T cells even when high affinity cognate antigen was present. This was unexpected given that during viral challenge, antigen encounter in peripheral tissue activates functional mechanisms of tissue retention in effector CD8^+^ T cells leading to the selective persistence of virus-specific CD8^+^ T cells and elimination of bystanders ^12^. Tumors, however, construct multiple biochemical and biophysical barriers that restrict T cell access to tumor nests and thus, to their target antigens ^9^, which may lead them to behave as bystanders. Consistent with this, we found that retained CD8^+^T cells were predominantly PD-1^+^ and progressively dysfunctional, indicating local antigen encounter, while egressed T cells were functional and PD-1^-^. These observations held even when evaluating clonal T cell populations with high affinity TCRs for exogenous neoantigens (e.g. OVA). Whether antigen-specific CD8^+^ T cells were retained or egressed depended on TCR-driven differential expression of chemokine receptors, notably CXCR4, ACKR3, and CXCR6 ^49, 50^, which identifies a causal link between antigen encounter in tumors, migration, and T cell function.

Similar to our results in CD8^+^ T cells, memory CD4^+^ T cells significantly downregulate surface CXCR4 in response to TCR stimulation *ex vivo* ^51, 52^ and exhibit impaired migration to CXCL12 ^53^. Interestingly, however, while antigen encounter drives CXCR4 surface loss, it also drives CXCR4 to associate with the TCR at the immune synapse in the presence of CXCL12 ^54^. This association increased IL-2, IL-4, and IL-10 production and enhanced AP-1 transcriptional activity (relative to NFAT and NF-κB activity) ^54^. This CXCR4-induced increase in AP-1 activity may have important implications as the balance of AP-1 and NFAT activity instruct T cell dysfunctional fates ^55^. Careful studies are needed to further determine the functional role of CXCR4-TCR crosstalk in effector CD8^+^ T cells and the consequences for T cell differentiation and dysfunction in tumors.

In addition to issues of antigen access, CD8^+^ T cell function in the TME is dictated by both antigen affinity and abundance ^37, 56, 57^. We found that the probability of T cell egress, as measured by CXCR4 and ACKR3 surface expression, was dependent on the strength of TCR stimulation, where low affinity antigen encounter (e.g., endogenous, self-antigens) was insufficient to promote retention. As such, TCR-induced retention mechanisms likely feed forward to promote chronic antigen exposure and exacerbate the accumulation of dysfunctional T cells in the TME. The TCF1^+^ stem-like T cell progenitor that gives rise to terminally dysfunctional effectors and is required for response to ICB ^31^, is enriched in clones with suboptimal TCR affinities in murine lung adenocarcinomas ^37^. This is consistent with our observation that TCF1^+^ stem-like and low affinity T cells egress from tumors, and may also indicate that afferent lymphatic transport helps to replenish TCF1^+^ stem-like T cells reservoirs in tumor dLNs for long term maintenance ^58^. Although more studies are needed to fully elucidate the complex interplay between antigen encounter, migration, and function, our findings support the hypothesis that tumors exploit physical barriers and weak antigens to expel a broad repertoire of functional CD8^+^ T cell clones.

Here, we present a model whereby local antigen encounter synergizes with the inflamed lymphatic vasculature to direct intratumoral CD8^+^ T cell repertoires. Our data indicates that lymphatic vessel-derived CXCL12 sequesters CD8^+^ T cells at the tumor periphery and facilitates their CXCR4-dependent egress from melanomas. Similarly, T cell exclusion in pancreatic cancers is CXCL12-dependent and genetic ablation of all peritumoral stromal fibroblasts (a subset of which produce CXCL12) improved T cell infiltration into tumor nests and enhanced spontaneous, T cell-dependent tumor control ^26^. In YUMMER1.7 melanoma models, only 2.6% (±2.1%) of tumor-associated fibroblasts expressed *Cxcl12* as compared to 8.9% (±4.8%) in naïve, dermal fibroblasts, and broad mapping of fibroblasts subsets indicate that *Cxcl12*^+^ activated fibroblasts are tissue-specific ^59^. Still, we do not exclude a role for fibroblast-produced CXCL12 in our model and it is interesting to consider how stromal and hematopoietic cell types might contribute competing chemokine gradients and diverse isoforms that ultimately shape the complex behavior or lymphocyte position in a tissue-specific manner.

Finally, consistent with our studies, CXCR4 inhibition improves intratumoral T cell accumulation and ICB efficacy in preclinical studies of pancreatic, breast, and hepatocellular cancer models ^26, 60, 61^. Due to the broad expression of CXCR4 in many cell types and tissues and the indiscriminate targeting by systemically administered inhibitors, it is difficult to tease out the exact mechanisms driving synergistic tumor control with ICB. Our study, however, indicates that these improvements may be due in part to impaired T cell egress, at least in models and patients with pre-existing antigen-specific immune responses. These preclinical studies importantly demonstrate the feasibility and potential impact of targeting lymphatic-mediated egress for combination immunotherapy. The tumor-associated lymphatic vasculature, and its transport function, is thereby an active player in regional disease control that may be leveraged to improve future patient outcomes.

## Methods

### Mice

C57Bl/6J, B6.SJL-*Ptprc^a^Pepc^b^*^/^BoyJ (CD45.1^+^), C57Bl/6-Tg(TcraTcrb)1100Mjb/J (OT1), B6.Cg(ROSA)26Sor^tm14(CAG-tdTomato)Hze^/J (tdTomato reporter line), B6(FVB-CXCL12^tm1.1Link^/J (CXCL12^fl/fl^), B6.129P2(Cg)-*Braf^tm1Mmcm^*/J (BRaf^V600E^), B6.129S4-*Pten^tm1Hwu^*/J (Pten^fl/fl^), and B6.Cg-Tg(Try-cre/ERT2)13Bos/J (Tyr::Cre-ER) were purchased from Jackson Laboratories. B6.Cg-Tg(CAG-tdKaede)-15Utr (Kaede-Tg) ^62^ were provided by D.J. Fowell in agreement with RIKEN BioResource Research Center; K14-VEGFR3-Ig ^21^ mice were provided by K. Alitalo (University of Helsinki, Finland); CXCL12-dsRedE2 reporter mice ^63^ were provided by I. Aifantis (NYU Langone) in agreement with S. Morrison (UT Southwestern); *Prox1:*Cre-ER^T2^ ^64^ were provided by V.H. Engelhard (University of Virginia) in agreement with T. Makinen (Uppsala University, Uppsala, Sweden); and CXCR4^fl/fl^-UBC-CreER^T2^ were provided by S.R. Schwab (NYU Langone; CXCR4^ΔUBC^). OT1 mice were crossed in house with Kaede-Tg to create OT1-Kaede-Tg mice. To specifically ablate CXCL12 from lymphatic endothelial cells (CXCL12^ΔiProx1^), CXCL12^fl/fl^ mice were crossed with mice expressing Cre recombinase behind the lymphatic-specific Prox1 promoter (*Prox1:*Cre-ER^T2^). Tamoxifen inducible lymphatic endothelial-specific tdTomato reporter mice (*Prox1*-tdTomato) were created by crossing *Prox1:*Cre-ER^T2^ with B6.Cg(ROSA)26Sor^tm14(CAG-tdTomato)Hze^/J mice. Tyr::Cre-ER, BRaf^V600E^, and Pten^fl/fl^ mice were crossed to generate the inducible autochthonous BPC melanoma model ^23^. For studies employing Kaede-Tg, K14-VEGFR3-Ig, CXCL12-dsRedE2, CXCL12^ΔiProx1^, and BPC mice, both male and female mice (8-20 weeks of age; age and sex matched) were utilized depending on availability. All experiments were performed in accordance with protocols approved by the Institutional Animal Care and Use Committees at New York University Langone Health and Oregon Health & Science University.

To induce systemic CreER^T2^-directed recombination, mice were treated with 2 mg tamoxifen (75 mg/kg; diluted in corn oil) for 5 consecutive days and then rested for at least 1 week. To induce localized cutaneous Tyr::Cre-ER recombination, ear skin of BPC mice were painted with 4-OH tamoxifen (7.1mg/ml); tumors developed 14-18 weeks post tamoxifen application.

For *in vivo* bromodeoxyuridine (BrdU) pulse studies, mice were intraperitoneally injected with 2 mg BrdU on day 0 of pulse followed by continual administration in the drinking water (0.8 mg/ml) for 4 days.

### Cell culture

YUMM1.7 and YUMMER1.7 were provided by M.W. Bosenberg (Yale University) and cultured as previously described ^65, 66^. MCA205 and MCA205.OVA were provided by J.M. Walker and A.E. Moran (Oregon Health & Science University) and cultured in DMEM supplemented with 10% FBS, 1% non-essential amino acids, 1mM sodium pyruvate, 10 mM HEPES, 1% penicillin/streptomycin. B16.F10 (ATCC CRL-6475), B16.F10.OVA (provided by M.A. Swartz at EPFL) and MC38 (provided by M.J. Gough at Providence Cancer Institute) tumor lines were cultured in DMEM supplemented with 10% FBS and 1% penicillin/streptomycin. All cell lines were maintained at 37°C with 5% CO_2_.

Primary lymph node (LN) stromal cells were cultured on tissue culture plates coated with 10 μg/ml PureCol (Sigma Aldrich) and 10 μg/ml fibronectin (Chemicon) in ⍺MEM (no nucleic acids) supplemented with 10% FBS and 1% penicillin and streptomycin. To induce CXCL12 gene expression ^46^, LN stromal cells were cultured under hypoxic conditions (1% O_2_) for 24 hours, and LN LECs sorted from cultures using fluorescence-activated cell sorting (FACs).

### In vivo tumor growth

For tumor implantation, 5×10^5^ tumor cells (suspended in 50μl saline) were injected intradermal using a 33-guage insulin syringe in the center of the upper back (drainage to both the left and right brachial LNs) or the left or right upper flank (drainage specifically to the left or right brachial LNs, respectively). Upon tumor palpation, the longest tumor diameter and diameter perpendicular to the longest diameter were measured with digital calipers. The formula 4/3νr^3^, where r is the one half the average of the longest diameter and the perpendicular diameter, was used to calculate tumor volume.

For CXCR4 inhibition, 10 mg/kg AMD3100 (Selleck Chemicals) was intraperitoneally administered daily. For immune checkpoint blockade, 200 μg of anti-mouse PD-L1 (clone 10F.9G2; Bio X Cell) antibodies or IgG isotype control were intraperitoneally administered every 3 days. Both AMD3100 and anti-PD-L1 therapies were initiated once tumor reached 200-250 mm^3^. For CD8 depletion, 200 μg anti-mouse CD8 (clone 2.43; Bio X Cell) antibodies were intraperitoneally administered twice per week starting the day of tumor implantation.

### *In vitro* and *in vivo* T cell activation

To create OVA-specific endogenous CD8^+^ effector T cells or to expand adoptively transferred OT1 CD8^+^ effector T cells, ActA-deficient OVA-expressing *Listeria monocytogenes* (LM-OVA) was grown in tryptic soy broth supplemented with 50 μg/ml streptomycin at 37°C with gentle shaking until 1×10^8^ CFU/ml (OD_600_=0.1). Following centrifugation, LM-OVA was resuspended in sterile PBS and 10^7^ CFUs (200 μl) intravenously transferred into mice. To activate OT-I CD8+ T cells, 5×10^4^ naïve OT1 TCR-Tg CD8^+^ T cell were intravenously transferred into naïve C57Bl/6J mice one day prior to infection. Spleens were harvested 7 days after infection and used for *in vit*ro studies. Single cell suspensions of spleens were generated by mechanical disruption through a 70um cell strainer. Red blood cells (RBCs) were lysed using VitaLyse (CytoMedical Design Group) per manufacturer’s instructions.

For *ex vivo* CD8^+^ T cell activation, OT-I splenocytes were stimulated with 1 nM SIINFEKL (OT-I CD8^+^ T cells) or in wells coated with anti-CD3 (10 μg/ml; clone: 145-2C11; Tonbo Biosciences) and anti-CD28 (2 μg/ml; clone: 37.51; eBiosciences) functional antibodies and 100 U/ml murine IL-2 for 72 hours. Activated CD8^+^ T cells were enriched using anti-APC microbeads (Miltenyi) and MACs LS Separation Columns (Miltenyi) per manufacturer’s instructions.

### Adoptive Transfer

0.5-4×10^6^ CD44^+^CD8^+^ T cells were intravenously transferred into tumor-bearing mice for adoptive transfer experiments. Alternatively, 1×10^6^ congenically labeled CD44^+^CD8^+^ T cells were intratumorally injected into mice bearing YUMMER1.7 tumors using a 33G Hamilton syringe (four 10 μl injections of 2.5×10^5^ splenocytes into four quadrants of the tumor).

### Kaede-Tg photoconversion

Kaede-Tg mice were photoconverted as previously described ^16^. Briefly, cutaneous tumors on anesthetized Kaede-Tg mice were exposed to 200 milliwatts of 405 nm light for 5 minutes, while surrounding skin was covered with aluminum foil to prevent photoconversion of non-tumor tissues. Tumors and corresponding tumor-draining LNs were harvested 24 hours post-photoconversion. In some cases, mice were treated with either pertussis toxin (1 μg/mouse) or FTY720 (25 μg/mouse) the day before and day of photoconversion.

### Tissue collection and processing

To create leukocyte single cell suspensions, LNs were teased open and digested with 1 mg/ml collagenase D and 80 U/ml DNase I in HBSS for 10-15 minutes at 37°C, then passed through a 70 μm cell strainer. Tumors were minced in DMEM containing 10% FBS and penicillin and streptomycin and then digested with 1mg/ml collagenase D and 80 U/ml DNase I in HBSS for 20-30 minutes with agitation. Tumors were then pressed through a metal screen followed by filtering through a 70 μm cell strainer. Splenocytes were pressed through a 70 μm cell strainer and then RBCs lysed with ACK buffer.

For isolation of cutaneous stromal cells, tumors and normal skin were minced and then digested with 220 U/ml collagenase IV (Invitrogen) and 80 U/ml DNase I at 37°C for 30 minutes with shaking. Tissue was pressed through a metal screen followed by filtering through a 70 μm cell strainer.

To isolate LN stromal cells (both for flow cytometry and cell culture), LN capsules were teased open with syringes and then digested in DMEM containing 220 U/ml collagenase IV and 80 U/ml DNase I for 30 minutes at 37°C with agitation. The supernatant was discarded and remaining stromal tissue washed twice with PBS to remove leukocytes. Stromal tissue was digested in DMEM containing 2% FBS, 3.3 mg/ml collagenase D, and 80 U/ml DNase I for 15 minutes at 37°C with agitation. Remaining tissue was then manually dissociated using a pipette ^67^.

Lymph was collected by thoracic duct cannulation using acid citrate dextrose-coated capillary tubes in lethally anesthetized mice. Blood collected from retroorbital sinus with heparin-coated capillary tubes followed by RBC lysis.

### Immunostaining for flow cytometry and fluorescence-activated cell sorting

Single cell suspensions were stained at 4°C in the dark with fixable live/dead dye diluted in PBS for 10 minutes followed by Fc-receptor blockade with Fc Shield (Tonbo Biosciences) for 15 minutes. For tetramer staining, single cell suspensions were incubated with H-2K^b^-OVA_257-264_- BV421 or H-2K^b^-gp_33-41_-APC for 45 minutes at room temperature in the dark. Tetramers were provided by the NIH Tetramer Core. For surface markers, single cell suspensions were incubated with primary antibodies diluted in PBS containing 1% BSA for 30-45 minutes at 4°C. For chemokine receptor staining, single cell suspensions were subsequently stained with chemokine receptor antibodies for 15 minutes at 37°C. Suspensions were then fixed with 2% PFA unless otherwise noted. For intracellular cytokine staining, single cell suspensions were fixed with BD CytoFix/CytoPerm (BD Biosciences) for 10 minutes at room temperature, followed by incubation with cytokine antibodies diluted in Perm Wash Solution (BD Biosciences) for 45 minutes at 4°C. Transcription factor staining was performed using True-Nuclear Transcription Factor Buffer Set (Biolegend) according to manufacturer’s instructions. For BrdU staining, single cell suspensions were fixed with CytoFix/CytoPerm for 30 minutes at 4°C followed by permeabilization with BD CytoPerm Plus Buffer (BD Biosciences) for 15 minutes at 4°C. Cell suspensions were then fixed again with CytoFix/CytoPerm for 10 minutes at room temperature followed by treatment with 0.33 mg/ml DNase I for 1.5 hours at 37°C. After washing with BD PermWash Buffer (BD Biosciences), cells were stained with anti-BrdU antibodies for 30 minutes at room temperature. For flow cytometry, samples were run through BD Fortessa or LSRII flow cytometers and data acquired using BD FACSDiva software. For FACS, cells were sorted using BD Influx, BD FACS Aria Fusion (OHSU Flow Cytometry Core), or a MoFlo-XDP flow sorters (NYU Langone Flow Cytometry and Cell Sorting Core).

**Table S6** lists antibodies and clones used for flow cytometry.

### *Ex vivo* restimulation and cytokine production

Day 21 YUMM1.7 or YUMMER1.7 tumors and draining LN single cell suspensions were restimulated for either 6 hours in 96-well plates coated with functional anti-mouse CD3 (10 μg/ml; clone: 145-2C11) and CD28 (2 μg/ml; clone: 37.51) antibodies or 3hrs with 500 ng/ml ionomycin and 250 ng/ml phorbol 12-myristate 13-acetate (PMA) in the presence of brefeldin A. For studies including CXCR4 blockade, 5 or 10 μg/ml of AMD3100 was added to the cultures.

### *Ex vivo* stimulation of OVA-specific effector CD8^+^ T cells with SIINFEKL peptides

Single cell suspensions were generated from the spleens of LM-OVA infected mice. Splenocytes were either incubated in plates coated with anti-CD3 and anti-CD28 functional antibodies or with 1 nM cognate peptides (SIINFEKL, SIIQFEKL, SIITFEKL) diluted in RPMI containing 10% FBS. In studies involving the addition of CXCL12, 100 ng/ml recombinant murine CXCL12 (R&D Systems) was added to the cultures. Cells were incubated at 37°C and 5% CO_2_ for times indicated in the figure.

### Effector CD8^+^ T cell transwell migration

Single cell suspensions were created from the spleens from LM-OVA infected mice. 1×10^6^ splenocytes resuspended in 100 μl serum free RPMI were add to the upper chamber of a 24-well transwell insert (Costar, pore size, 5.0 μm). 600 μl of either serum free media (control) or 100 ng/ml recombinant murine CXCL12 was added to the lower well. For experiments requiring peptide stimulation, 1 nM SIINFEKL was included with the splenocytes in the upper chamber. Cells were incubated at 37°C and 5% CO_2_ for 4 hours. After incubation cells in the lower well were collected and analyzed by flow cytometry.

### TCR deep sequencing

Genomic DNA was extracted with the QIAamp DNA isolation kit (Qiagen) per manufacturer’s instruction and then submitted for TCR deep sequencing. TCR deep sequencing library was prepared using a two-stage PCR procedure as previously described ^68^. Briefly, genomic DNA and synthetic TCR internal control templates were amplified in a first stage multiplex PCR by 20 V forward and 13 J reverse primers using the Qiagen multiplex PCR kit. Using 2% of the purified first-PCR product, a second stage PCR was performed to add universal and index Illumina adaptors. Equal volumes of each sample were pooled to create the TCR sequencing library that was profiled using a 2200 TapeStation (Agilent) and quantified by qRT-PCR (Thermo Fisher Scientific’s StepOne Real Time Workstation) with the Kapa Biosystems commercial library quantification kit. The library was then sequenced with a 2 x 150 protocol on Illumina NextSeq 500 platform using a Midoutput 300 sequencing kit at 160 million target clusters per run (OHSU Massively Parallel Sequencing Shared Resource Core). In-house data analysis pipeline along with MiXCR ^69^ and tCR ^70^ packages were used to process the data, identify purified clone sequences, and normalize clone counts. TCR repertoire metrics were calculated as previously described ^68^.

### Bulk RNA-Seq

To compare egressed and tumor retained CD8^+^ T cell transcriptomes, single cell suspensions were generated from YUMM1.7 tumors and dLNs as described above. Cells were immunostained as described above and live (DAPI^-^) Kaede red^+^CD44^+^CD8^+^ T cells were sorted into RPMI containing 10% FBS using a BD FACS Aria sorter. Cells were lysed using Takara Bio Single Cell Lysis Buffer containing a recombinant RNase inhibitor (Takara Bio Inc.) then snap frozen. Bulk RNA-sequencing and sequence alignment was performed by MedGenome Inc (Foster City, CA, USA). Complementary DNA libraries were prepared using Takara SMART-Seq v4 Ultra low input RNA kit and quality assessed using Qubit Fluorometric Quantitation and TapeStation bioanalyzer. Products were sequenced using an Illumina NovaSeq platform with a target of 40 million unpaired reads per sample. Sequence alignment was performed using STAR aligner (v2.7.3a) and raw counts estimated using HTSeq (v.0.11.2). From aligned reads, normalized counts were generated using DESeq2 and estimated gene expression generated using cufflinks (v2.2.1).

To evaluate the transcriptional differences between naïve and tumor-associated LECs, cutaneous BPC tumors (induced on ear pinna) and naïve ear skin were harvested from mice and the dorsal and ventral sides of the pinna separated from one another. To create single cell suspensions both halves of the pinna were enzymatically digested with 220 U/ml collagenase IV and 80 U/ml DNase I at 37°C for 2hrs while shaking. Tissues were then pressed through a metal screen followed by filtering through a 70μm cell strainer. Cells were immunostained with anti-mouse CD31 (MEC13.3), gp38 (8.1.1) and CD45 (30-F11) antibodies as described above. LECs (CD45^-^ CD31^+^gp38^+^) were sorted using a BD Influx flow sorter (OHSU Flow Cytometry Core). Sorted LECs were submitted to OHSU’s Gene Profiling Shared Resource Core for RNA Isolation using RNeasy Micro Kit (Qiagen); RNA quality was assessed using Bioanalyzer 2100 with RNA 6000 Pico chips. Products were sequences using an Illumina HiSeq 2500 platform (OHSU Massively Parallel Sequencing Shared Resource). Sequence alignment was performed using STAR aligner (v2.6). DESeq2 ^71^ was used to identify differentially expressed genes; results were transformed and scaled z-scores plotted in heatmaps generated using pheatmap or ComplexHeatmap R packages. Gene set enrichment was performed using fgsea and Msigdbr R packages.

### Human melanoma scRNA-Seq

The existing scRNA-Seq dataset GSE120575 collected from human melanomas (both untreated and immune checkpoint blockade treated) was obtained from the National Center for Biotechnology Information ^34^. Single cell counts were loaded into R and data processing, visualization, and analysis performed using Seurat (v4) R package ^72^. Data were normalized and scaled using the ‘sctransform’ function in R. FindNeighbors and FindCluster Seurat functions were used to identify *CD3^+^CD8^+^CD4^-^*T cell clusters and Uniform Manifold Approximation and Projection (UMAP) was used for visualization. DESeq2 was used to determine differential gene expression among identified cell clusters.

### RNAScope *in situ* hybridization

Formalin-fixed paraffin-embedded 5 μm murine tumor sections were baked overnight at 60°C. Tissue dehydration and RNA *in situ* hybridization was performed according to manufacturer’s instructions using RNAScope 2.5 HD Assay-RED kit and manufacturer-designed probe Mm-*Cxcl12* (422711) from Advanced Cell Diagnostics.

### Immunofluorescence microscopy

Tumors were fixed with 1% PFA for 24-48 hours at 4°C, followed by sequential incubations in 15% sucrose and 30% sucrose for 24 hours each at 4°C. Tumors were then embedded in OCT and cut into 8 μm sections. Tissue sections were blocked with 2.5% BSA for 30-45 minutes at room temperature. For CXCL12 immunostaining, tissue sections with treated with Mouse-on-Mouse Immunodetection Kit reagents (Vector Laboratories) according to manufacturer’s instructions followed by staining with additional primary antibodies (see **table S6** for antibody list). After blocking, unconjugated primary antibodies (diluted in 1.25% BSA) were incubated overnight at 4°C. Sections were then washed once with TBS+0.1% Tween-20 (TBST) and twice with PBS. Fluorophore-conjugated primary and secondary antibodies (diluted in 1.25% BSA) were then applied to the tissue sections and incubated for 2 hours in the dark at room temperature. Sections were again washed with TBST and PBS. After secondary antibody incubations, sections were treated for 10 minutes with DAPI nuclear stain (Life Technologies), washed and then mounted using SlowFade Gold antifade reagent (Invitrogen). Fluorescence images were captured using Keyence BZ-X810 All-In-One Fluorescence Microscope. For quantification, single-channel images were converted to 8-bit format and positive pixels per image enumerated using Fiji opensource software (National Institutes of Health).

### Multiplex immunohistochemistry (mIHC) on primary human melanomas

Deidentified 5 μm formalin fixed paraffin-embedded primary human melanoma (stage I-III) tissue sections were acquired from the OHSU Knight Biolibrary and OHSU Department of Dermatology research repository in accordance with OHSU’s Institutional Review Board. In total 28 patient melanoma samples (14 male and 14 female) were stained and analyzed. Assessment by a dermatopathologist confirmed the presence of melanoma in each sample. Average tumor thickness was 2.86 (±1.68) mm; 11 patients had positive LN biopsies while 17 patients had negative LN biopsies.

Using a modified protocol for melanoma samples, sequential chromogenic immunohistochemistry was performed on each primary melanoma tissue section ^73^. Briefly, tissue sections were deparaffinized then bleached with 10% H_2_O_2_ at 65°C for 10 minutes. After bleaching, sections were stained with hematoxylin for 1 minute followed by whole slide imaging using an Aperio ImageScope AT (Leica Biosystems) at 20X magnification. After scanning, heat-induced epitope retrieval with Citra Plus solution (BioGenex) was performed for 20 minutes. Subsequent iterative cycles of standard immunohistochemistry were performed using the following anti-human primary antibodies: podoplanin (D240, BioLegend; 1:100), LYVE-1 (rabbit polyclonal, Abcam; 1:100), αSMA (rabbit polyclonal, Abcam; 1:500), S100 (clonal cocktail, Biocare Medical; 1:2000), CD8 (C8/144B, Thermo Fisher; 1:200), CD34 (QBEnd-10; Thermo Fisher; 1:10000), aquaporin 1 (rabbit polyclonal, Millipore; 1:10000), and panCK (AE1/AE3, Abcam; 1:2000). Species-specific ImmPress horseradish peroxidase polymer reagent and AEC (Vector Laboratories) were used for antigen detection and visualization. After whole slide scanning of each cycle, AEC was removed using ethanol and antibodies stripped using heated Citra Plus solution. Hematoxylin staining was repeated after the final staining cycle.

### mIHC image processing and analysis

Processing, image alignment and registration, and pseudo-color visualization of serial digitized images were performed as previously described ^73^. For analysis, three rectangular 6.25 mm^2^ regions of interest (ROIs) were selected per section based on 1) presence of podoplanin^+^ lymphatic vessels and 2) inclusion of stromal tissue and tumor parenchyma. To differentiate between stroma and tumor parenchyma in the ROIs, tumor segmentation masks were generated as previously described ^7^. This was done by first triangle thresholding of hematoxylin-stained nuclei (for tissue detection) followed by S100^+^ detection via computing an alternate sequential filter and triangle thresholding. An inverted mask was used to segment peritumoral tissue. These masks were used to identify CD8^+^ T cell as peritumoral or intratumoral; these cells were quantified using CellProfiler (v3.5.1).

Vessel segmentation was performed using Otsu’s method to segment blood and lymphatic vessels based on AQP1^+^, CD34^+^ (blood) and podoplanin^+^ (lymphatic) staining; panCK staining was used to excluded podoplanin^+^ basal epithelial cells. Mathematical morphology operation was used to denoise and locally flatten single channels images, which were then used in Otsu’s thresholding to create a binarized images of lymphatic vessels structures. Size-based filtering was applied to remove staining artifacts. Vessel subtypes (LYVE-1^+^ vs LYVE-1^-^) were identified using supervised hierarchical gating of positive pixel coverage and manual intensity thresholding of vessel markers.

### RNA isolation and qRT-PCR

Total RNA was isolated fro using RNAqueous Micro Total RNA Isolation Kit (Thermo Fisher Scientific) according to manufacturer’s instructions. 100 ng of RNA was reverse transcribed into cDNA using High-Capacity cDNA Reverse Transcription Kit (Thermo Fisher Scientific). cDNA was then diluted 1:10 and qPCR was performed using PowerUp SYBR Green Master Mix (Thermo Fisher) and murine CXCL12 or β-actin forward and reverse primers in a 384-well plate format. Primers were designed using NCBI-BLAST: CXCL12-forward primer 5’-CCTTCAGATTGTTGCACGGC-3’; CXCL12-reverse primer 5’-CTTGCATCTCCCACGGATGT-3’; β-actin-forward primer 5’-CTGTCCCTGTATGCCTCTG-3’; β-actin-reverse primer 5’-ATGTCACGCACGATTTCC-3’. A BioRad CFX384 Touch Real-Time PCR Detection System was used for thermocycling and signal detection. CXCL12 expression was normalized based on β-actin expression and relative expression calculated using 2^-ΔΔCT^.

### Statistical analysis

Statistical analyses were performed using GraphPad Prism (v9) software using parametric or non-parametric Student’s t-tests and one- and two-way ANOVA for multiple pairwise testing as indicated. Asterisks indicate significance level: *p<0.05, **p<0.01, ***p<0.001, and ****p<0.0001. n.s. = not significant. Specific statistical tests performed for each experiment are listed in the figure legends.

## Acknowledgments

The authors would like to acknowledge Susan R. Schwab for critical input, and members of the Lund Lab for critical feedback and technical support.

## Funding

National Institutes of Health grant P30-CA0695533 (LMC, AWL)

National Institutes of Health grant R01CA238163 (AWL)

National Institutes of Health grant T32CA106195 (MMS)

National Institutes of Health grant T32GM136542 (HdB)

Cancer Research Institute, Lloyd J. Old STAR Award (AWL)

Melanoma Research Alliance 403191; https://doi.org/10.48050/pc.gr.44893 (AWL)

Brenden-Colson Center for Pancreatic Health (LMC)

Stand Up to Cancer – Lustgarten Foundation Pancreatic Cancer Convergence Dream Team Translational Research Grant SU2C-AACR-DT14-14 (LMC)

Swedish Research Council (Vetenskapsrådet), International Postdoc grant, Reg.Nr:2016-00215 (JF)

National Institutes of Health grant P30-CA016087 (Laura and Isaac Perlmutter Cancer Center supporting the Flow Cytometry and Cell Sorting Core and the Experimental Pathology Research Laboratory)

National Institutes of Health grant P30-CA069533 (OHSU Knight Cancer Center supporting the Flow Cytometry and Cell Sorting Core, Massively Parallel Sequencing Shared Resource, and Gene Profiling Shared Resource)

## Author contributions

Conceptualization: MMS, AWL

Methodology: MMS, CH, DM, JF, YHC

Biospecimen Acquisition: SAL

Investigation: MMS, IDD, HdB, DM, JF

Funding acquisition: LMC, AWL

Supervision: LMC, AWL

Writing – original draft: MMS, AWL

Writing – review & editing: MMS, IDD, HdB, DM, CH, JF, LMC, AWL

## Data and materials availability

All sequence data, code, and materials used in the analysis will be made publicly available upon manuscript acceptance.

---

**Supplementary Fig. 1.**
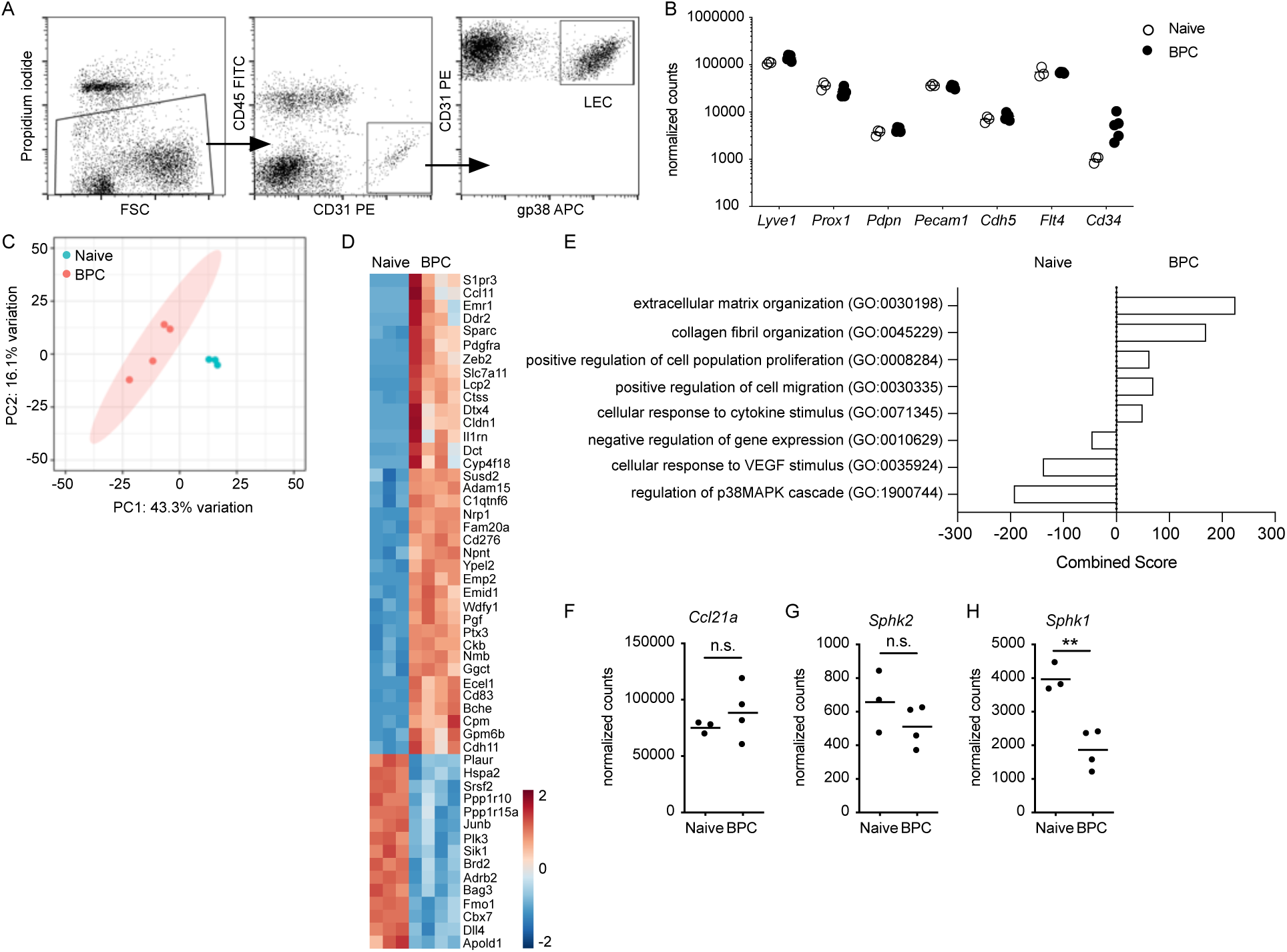
RNA-Seq of LECs sorted from naïve skin and BPC melanomas. (**A**) Gating scheme for FACS of LECs (CD45^-^CD31^+^gp38^+^) from naïve skin and BPC tumors. (**B**) Normalized transcript counts of genes validating LEC identity for CD45^-^CD31^+^gp38^+^ cells sorted from naïve skin and BPC tumors. (**C**) Principal component analysis comparing the transcriptomes of LECs sorted from naïve skin and BPC tumors. (**D**) Heatmap of top differentially expressed genes among naïve and BPC-associated LECs. (**E**) Gene set enrichment analysis of pathways enriched in naïve vs BPC-associated LECs. (**F-H**) Normalized transcript counts for *Ccl21a* (F), *Sphk2* (G), and *Sphk1* (H) in naïve and BPC-associated LECs. Each symbol represents one mouse; n=3 naïve and 4 BPC mice. **p<0.01, n.s.= not significant

**Supplementary Fig. 2.**
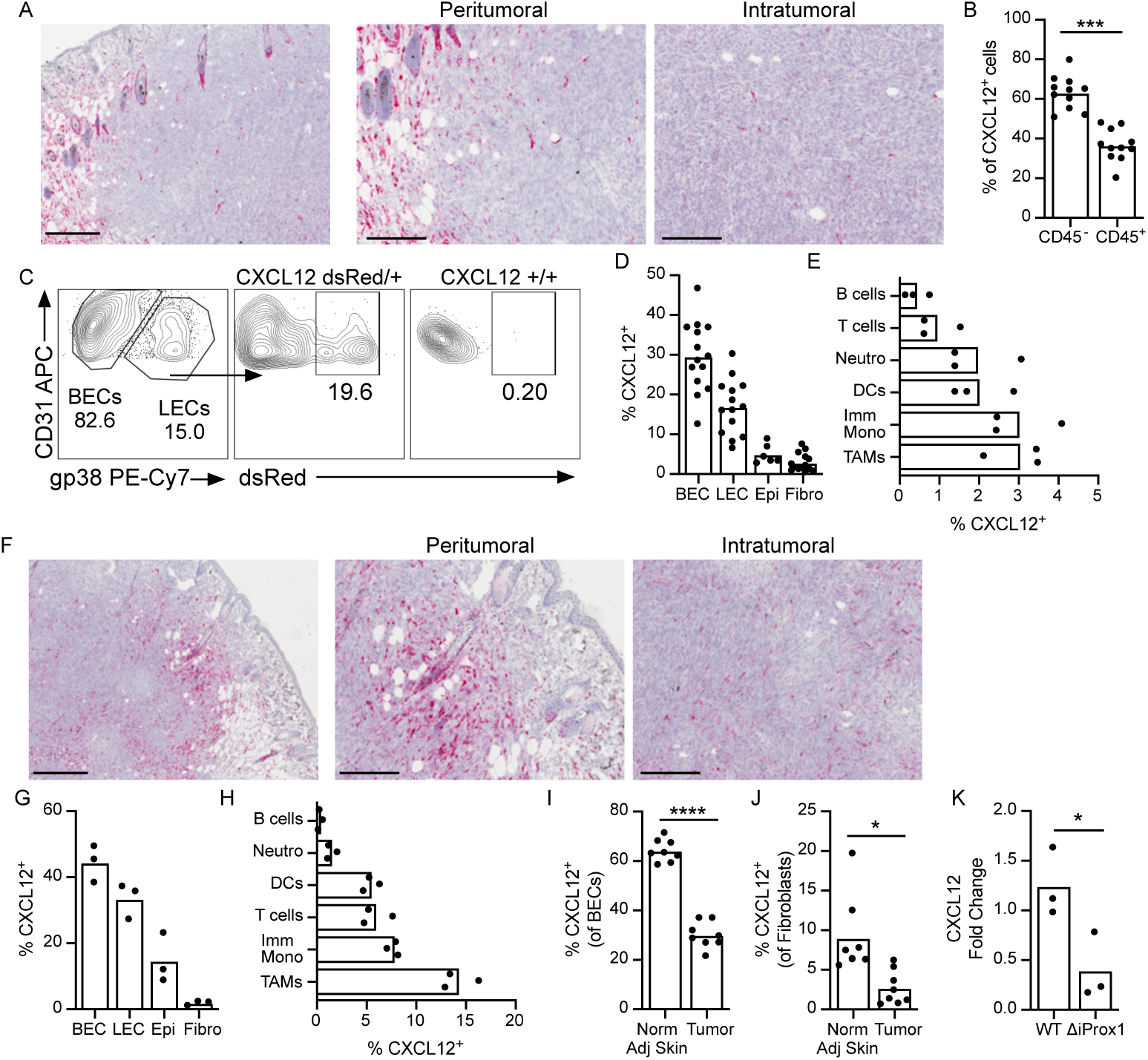
Expression of *Cxcl12* transcripts in YUMMER1.7 and YUMM1.7 tumor microenvironments. (**A**) RNAScope *in situ* staining of *Cxcl12* transcripts in YUMMER1.7 tumors; counterstained with hematoxylin. Repeated in 3 tumors. Left image: scale bar = 400 μm; middle and right images: scale bar = 200 μm (**B**) Frequency of *Cxcl12*-dsRed^+^ cells that are either CD45^+^ or CD45^-^ in YUMMER1.7 tumors implanted in *Cxcl12*-dsRed reporter mice (n=11 across 2 independent experiments. (**C**) Representative flow plots identifying CD31^+^gp38^+^dsRed^+^ LECs in *Cxcl12*-dsRed reporter mice or littermate controls (*Cxcl12*^+/+^). (**D**) Frequency of *Cxcl12*^+^ cells among CD45^-^CD31^+^gp38^-^ blood endothelial cells (BECs), CD45^-^CD31^+^gp38^+^ LECs, CD45^-^CD31^-^EpCAM^+^ epithelial cells (Epi), CD45^-^CD31^-^ PDGFR ^+^ fibroblasts (Fibro) in YUMMER1.7 tumors of *Cxcl12*-dsRed reporter mice (n=13; 3 independent experiments). (**E**) Frequency of *Cxcl12*^+^ cells among CD3 ^-^B220^+^ B cells, CD3 ^+^B220^-^ T cells, CD3 ^-^B220^-^ CD11b^+^Ly6G^+^ neutrophils (Neutro), CD3 ^-^B220^-^CD11c^+^MHCII^+^ dendritic cells (DCs) CD3 ^-^B220^-^ CD11b^+^Ly6C^+^ immature monocytes (Imm Mono), or CD3 ^-^B220^-^CD11b^+^Ly6C^-^F4/80^+^MHCII^+^ macrophages (TAMs) in YUMMER1.7 tumors of *Cxcl12*-dsRed reporter mice (n=3). (**F**) RNAScope *in situ* staining of *Cxcl12* transcripts in YUMM1.7 tumors; counterstained with hematoxylin. Repeated in 3 tumors. Left image: scale bar = 400 μm; middle and right images: scale bar = 200 μm (**G**) Frequency of *Cxcl12*^+^ cells among CD45^-^CD31^+^gp38^-^ BECs, CD45^-^CD31^+^gp38^+^ LECs, CD45^-^CD31^-^EpCAM^+^ epithelial cells (Epi), CD45^-^CD31^-^gp38^+^PDGFR ^+^ fibroblasts (Fibro) in YUMM1.7 tumors of *Cxcl12*-dsRed reporter mice (n=3). (**H**) Frequency of *Cxcl12*^+^ cells among CD3 ^-^B220^+^ B cells, CD3 ^+^B220^-^ T cells, CD3 ^-^B220^-^ CD11b^+^Ly6G^+^ neutrophils (Neutro), CD3 ^-^B220^-^CD11c^+^MHCII^+^ DCs, CD3 ^-^B220^-^CD11b^+^Ly6C^+^ immature monocytes (Imm Mono), or CD3 ^-^B220^-^CD11b^+^Ly6C^-^F4/80^+^MHCII^+^ macrophages (TAMs) in YUMM1.7 tumors of *Cxcl12*-dsRed reporter mice (n=3). (**I-J**) Frequency of *Cxcl12*^+^ cells among CD45^-^ CD31^+^gp38^-^ BECs (I) or CD45^-^CD31^-^PDGFR ^+^ fibroblasts (J) from normal adjacent skin or YUMMER1.7 tumors in *Cxcl12*-dsRed reporter mice (n=8; 2 independent experiments). (**K**) qRT-PCR of relative CXCL12 transcript levels from hypoxic LN LECs isolated from CXCL12^WT^ or CXCL12^ΔiProx1^ mice (n=3). Hypoxia was used to induce CXCL12 expression in primary LEC cultures; LECs were sorted from contaminating fibroblast reticular cells based on CD31 expression. For all graphs, each symbol represents one mouse. Two-sided paired (B, I, and J) and unpaired (K) student’s t-test. *p<0.05, ***p<0.001, ****p<0.0001.

**Supplementary Fig. 3.**
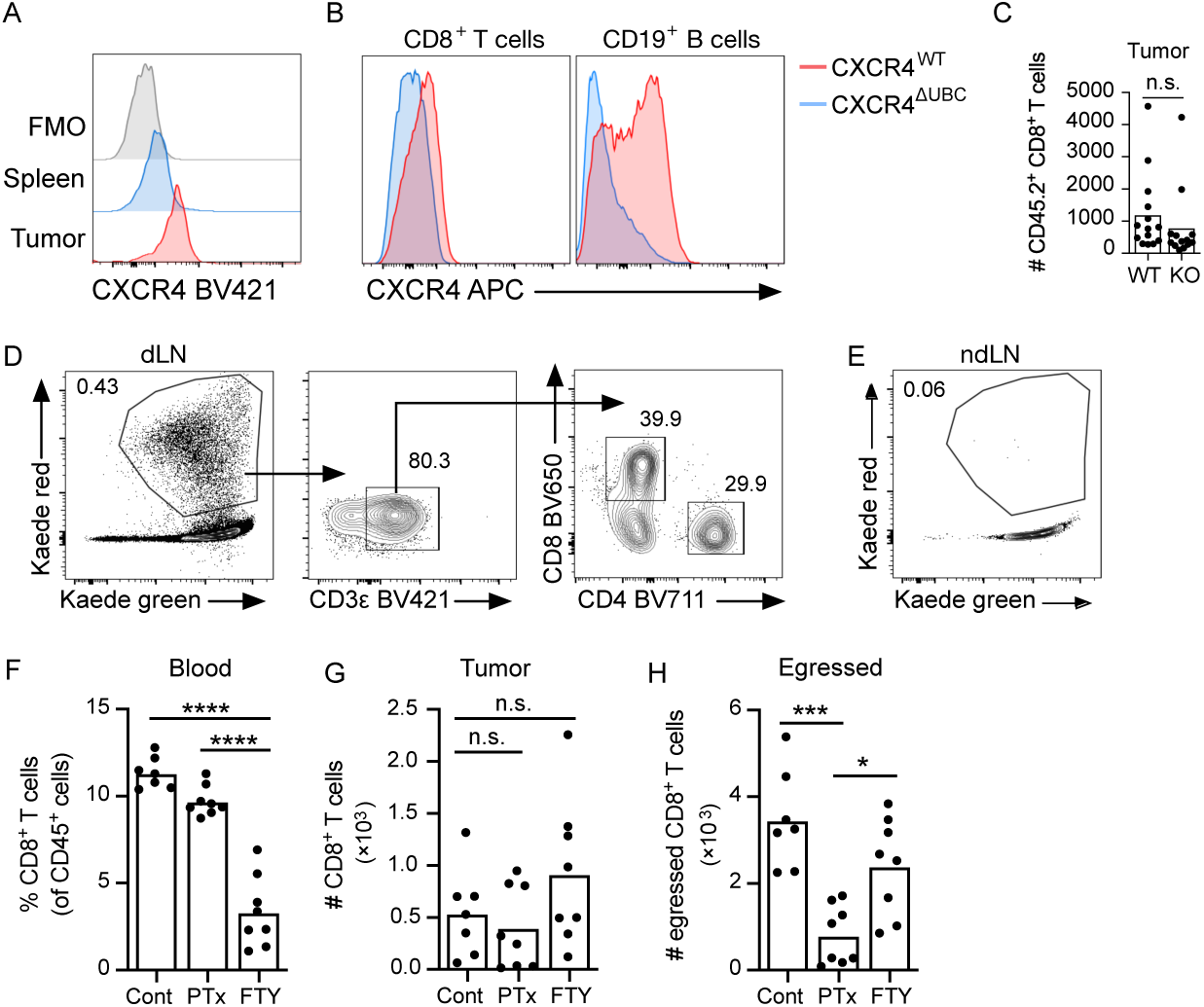
CD44^+^CD8^+^ T cells egress from tumors is G protein-coupled receptor and CXCR4-dependent. (**A**) Representative histogram of CXCR4 surface expression on CD44^+^CD8^+^ T cells from spleen and YUMMER1.7 tumors. FMO = Fluorescence minus one (CXCR4). (**B**) Histograms confirming CXCR4 knockout in CD8^+^ T cells (left panel) or CD19^+^ B cells (right panel) from CXCR4^WT^ (red) or CXCR4^ΔUBC^ (blue) mice. (**C**) Numbers of CD45.2^+^CD8^+^ CXCR4^WT^ (WT; n=14)) or CXCR4^ΔUBC^ (KO; n=13) T cells in YUMMER1.7 tumors 16-24hrs post intratumoral transfer (1×10^6^ CD8^+^ T cells). Each symbol represents one mouse. (**D**) Representative gating scheme identifying egressed Kaede red^+^CD8^+^ T cells in tumor draining brachial LNs 24hrs post photoconversion of the tumor. (**E**) Representative flow plot demonstrating no photoconverted cells in non-draining inguinal LNs 24hrs post photoconversion of the tumor. (**F-H**) YUMM1.7-bearing Kaede-Tg mice were treated with FTY720 (FTY) or pertussis toxin (PTx) the day before and day of photoconversion. (F) Frequency of CD8^+^ T cells (among live CD45^+^ cells) in the blood post treatment with pertussis toxin (PTx) or FTY720 (FTY). (G) Numbers of CD8^+^ T cells in YUMM1.7 tumors of Kaede-Tg mice post treatment with PTx or FTY720. (H) Numbers of Kaede red^+^CD8^+^ T cells in the dLNs of YUMM1.7 tumors of Kaede-Tg mice post treatment with PTx or FTY720. For F-H, n= 7 vehicle control mice; n=8 PTx-treated mice; n=8 FTY-treated mice; each symbol represents one mouse; results of 2 independent experiments. Two-sided, unpaired student’s t-test (C) and one-way ANOVA (F-H). *p<0.05, ***p<0.001, ****p<0.0001, n.s. = not significant.

**Supplementary Fig. 4.**
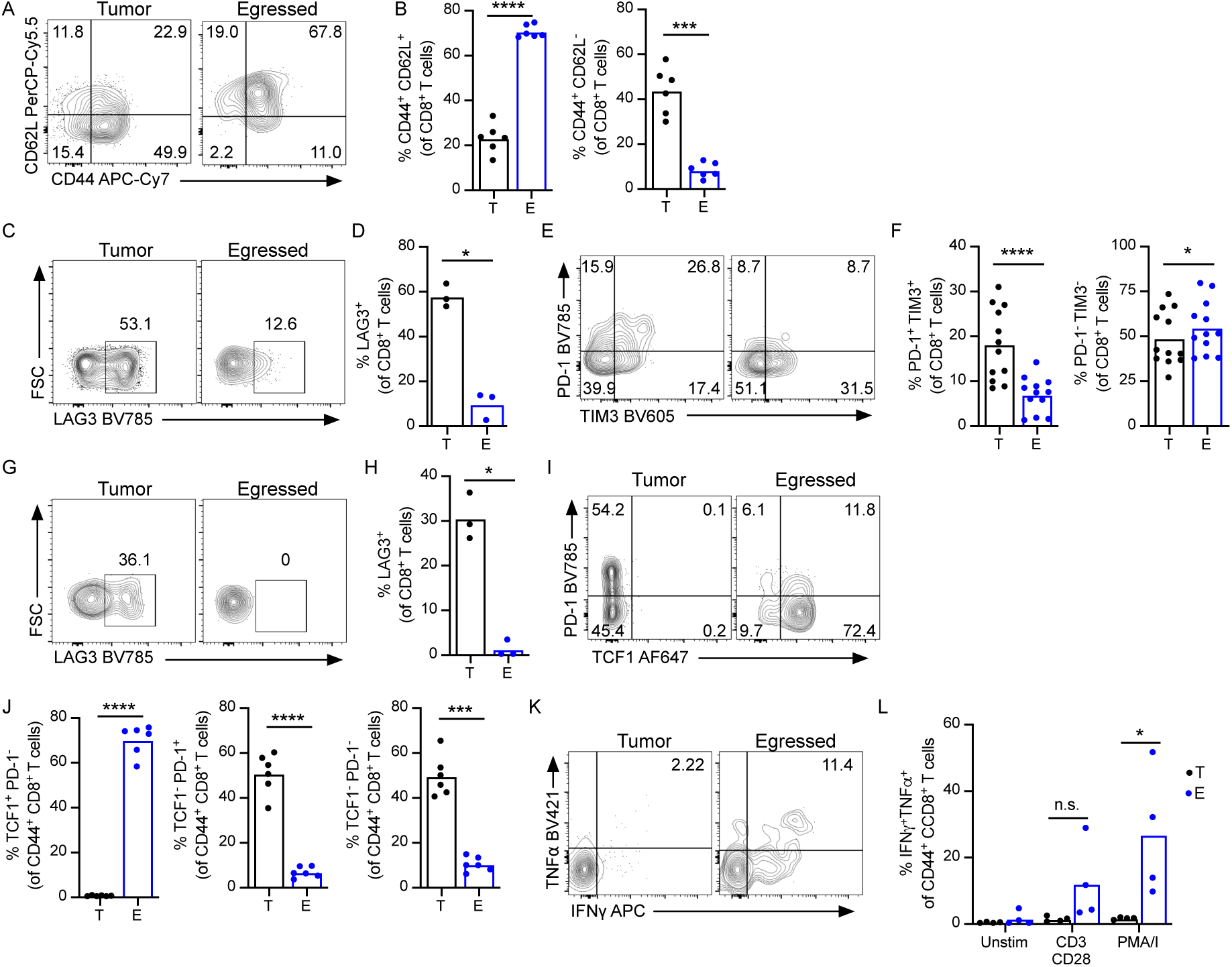
Functional effector CD44^+^CD8^+^ T cells egress from melanoma microenvironments. (**A-B**) Representative flow plots (A) and frequencies (B) of CD44/CD62L cell populations among intratumoral (T) or egressed (E) CD8^+^ T cells from YUMMER1.7-bearing Kaede-Tg mice (n=6) 24hrs post photoconversion. (**C-D**) Representative flow plots (C) and frequencies (D) of LAG3^+^ cells among intratumoral (T) or egressed (E) CD44^+^CD8^+^ T cells from YUMMER1.7-bearing Kaede-Tg mice 24hrs post photoconversion. (**E-F**) Representative flow plots (E) and frequencies (F) of PD-1/TIM3 cell populations among intratumoral (T) or egressed (E) CD44^+^CD8^+^ T cells from YUMM1.7-bearing Kaede-Tg mice (n=12; from 3 independent experiments) 24hrs post photoconversion. (**G-H**) Representative flow plot (G) and frequencies (H) of LAG3^+^ cells among intratumoral (T) or egressed (E) CD44^+^CD8^+^ T cells from YUMM1.7-bearing Kaede-Tg mice (n=3) 24hrs post photoconversion. (**I-J**) Representative flow plots (I) and frequencies (J) of PD-1/TCF1 cell populations among intratumoral (T) or egressed (E) CD44^+^CD8^+^ T cells YUMMER1.7-bearing Kaede-Tg mice (n=6) 24hrs post photoconversion. (**K-L**) Representative flow plots (K) and frequency (L) of *ex vivo* IFN and TNF production by intratumoral (T) or egressed (E) CD44^+^CD8^+^ T cells from YUMM1.7 tumors (day 21; n=4) following stimulation with functional CD3/C28 antibodies or PMA and ionomycin (PMA/I). For all graphs, each symbol represents one mouse. Two-sided, paired and (B, D, F, H, and J) and unpaired student’s t-test (L). *p<0.05, ***p<0.001, ****p<0.0001, n.s. = not significant.

**Supplementary Fig. 5.**
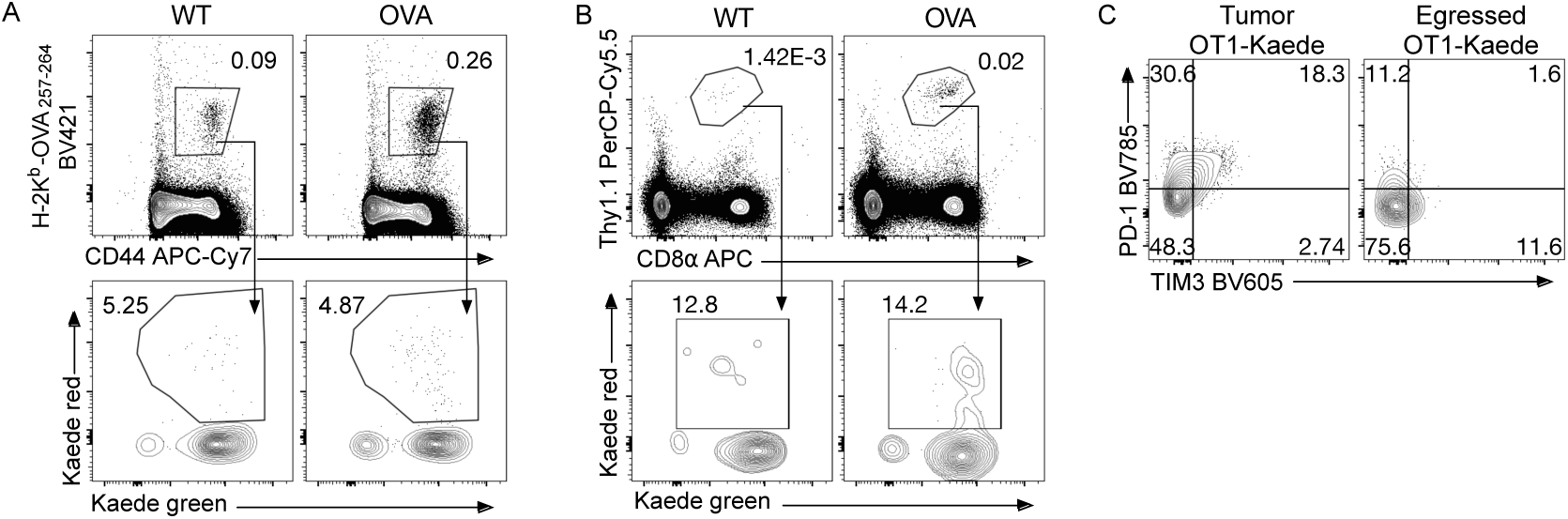
Identifying egressed tumor-specific CD44^+^CD8^+^ T cells in tumor draining LNs. (**A**) Representative gating scheme identifying endogenous egressed Kaede red^+^ H-2K^b^-OVA_257-264_^+^ CD44^+^CD8^+^ T cells in tumor draining brachial LNs 24hrs post photoconversion of MCA205^WT^ (WT) or MCA205^OVA^ (OVA) tumors. (**B**) Representative gating scheme identifying egressed Kaede red^+^ Thy^1.1/1.1^CD44^+^CD8^+^ OT1 T cells in tumor draining brachial LNs 24hrs post photoconversion of B16.F10^WT^ (WT) or B16.F10^OVA^ (OVA) tumors. (**C**) Representative flow plots of PD-1/TIM3 cell populations among intratumoral (T) or egressed (E) CD44^+^CD8^+^ OT1-Kaede-Tg T cells from B16.F10^OVA^-bearing mice 24hrs post photoconversion.

**Supplementary Fig. 6.**
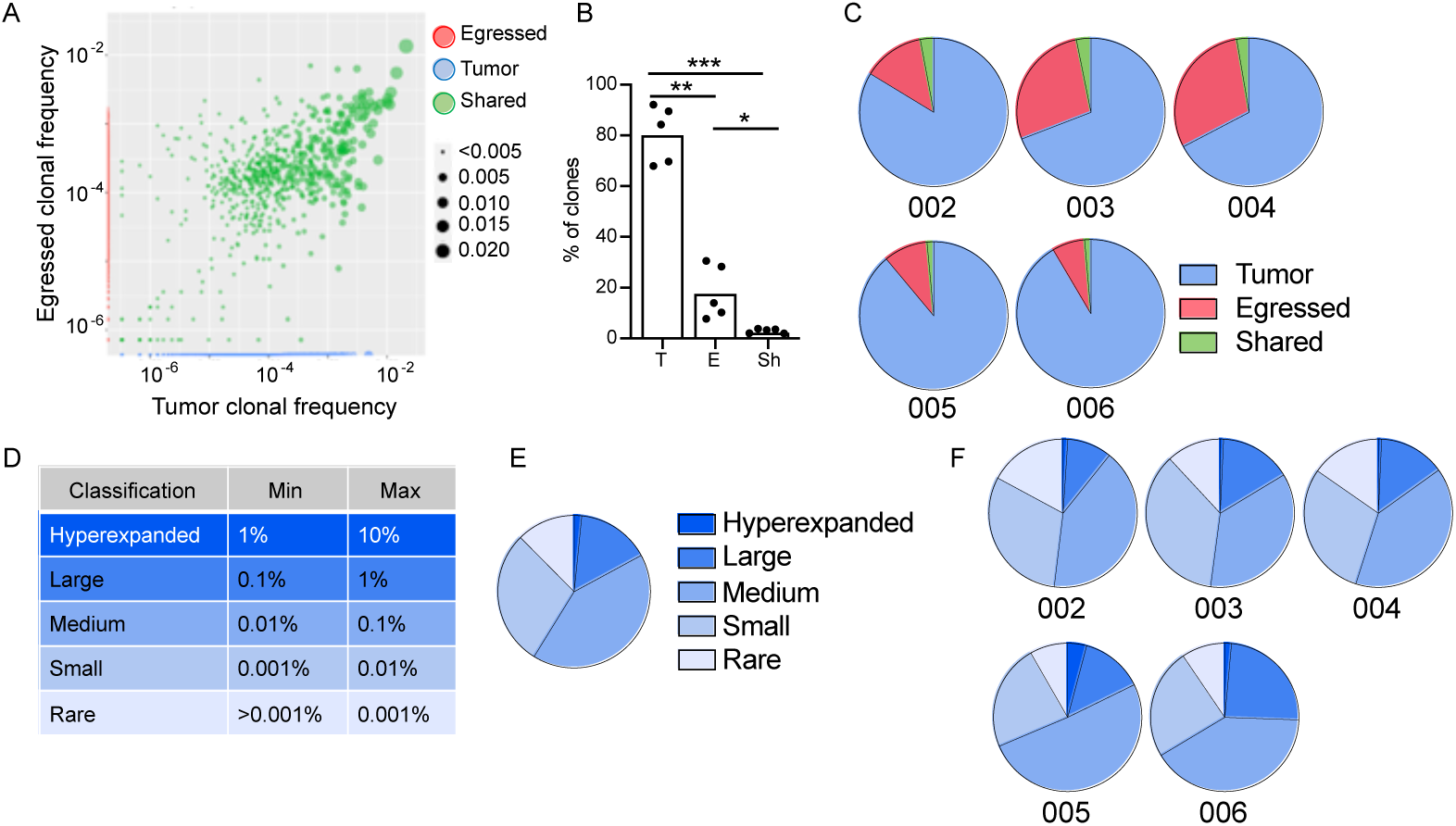
TCR-Sequencing of intratumoral and egressed T cells. Kaede red^+^ CD3 ^+^ T cells were sorted from YUMM1.7 draining LNs 24hrs post photoconversion and submitted for TCR deep sequencing along with matched YUMM1.7 T cells (n=5). In total, 5 mice were submitted for sequencing (**A**) Representative bubble plot (mouse 003) identifying clones sequenced in the egressed compartment (red), tumor compartment (blue), or both (Shared; green). (**B**) The percentage of sequenced clones detected in the tumor (T) or egressed (E) compartments, or both (shared; Sh). Each symbol represents one mouse. One-way ANOVA was used to determine statistical significance. *p<0.05, **p<0.01, p<0.001. (**C**) Pie charts for each mouse indicating the percentage of sequenced clones detected in the tumor (blue) or egressed (red) compartments or both (Shared; green). (**D**) Table demonstrating the clonal size criteria for classifying clones as hyperexpanded, large, medium, small or rare clones. (**E**) Pie chart demonstrating the percentage of shared clones classified as hyperexpanded, large, medium, small or rare clones averaged across all mice. (**F**) Pie charts demonstrating the percentage of shared clones classified as hyperexpanded, large, medium, small or rare clones for each individual mouse.

**Supplementary Fig. 7.**
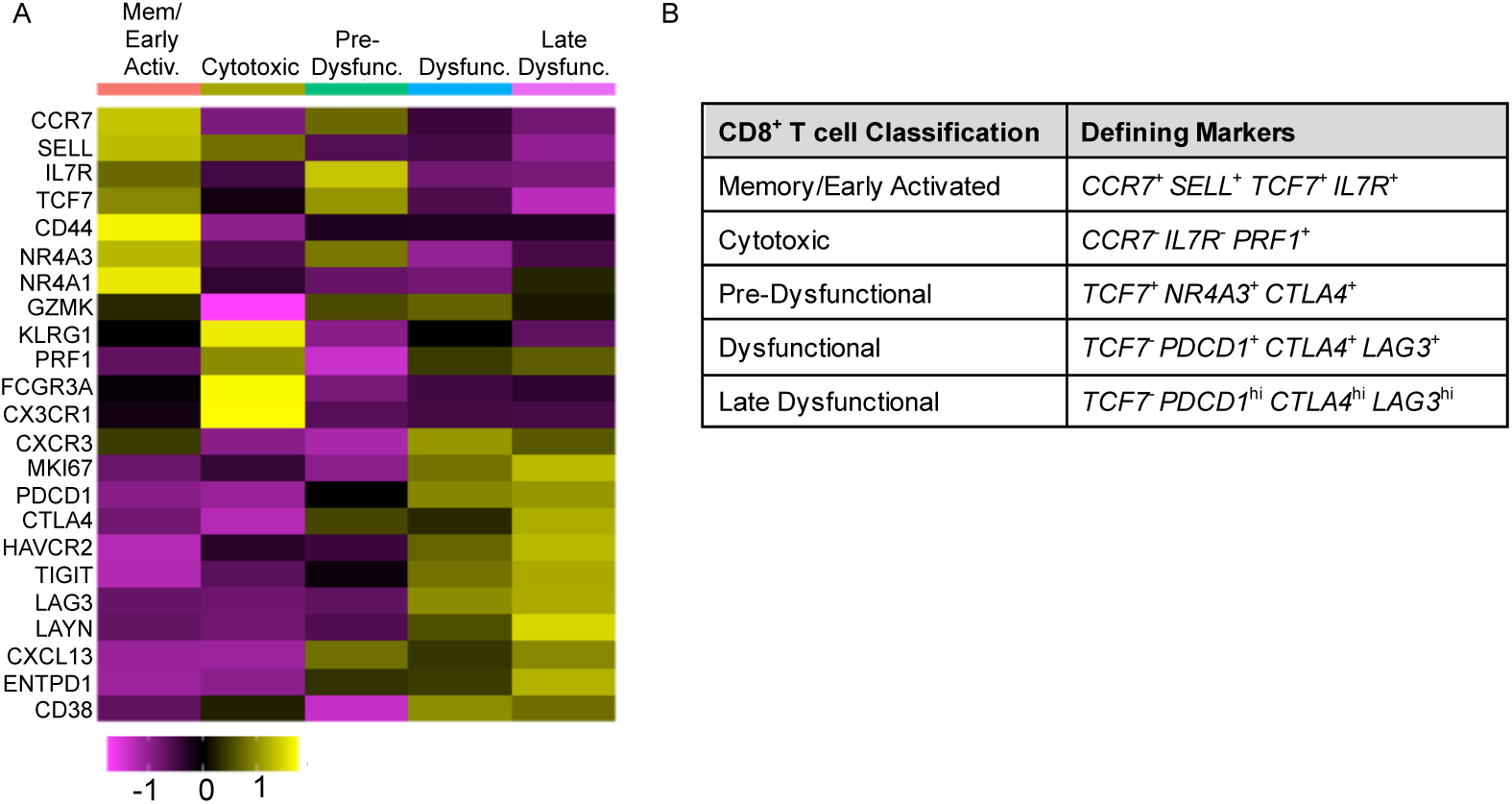
CD8^+^ T cell functional states in human melanomas. Identification of CD8^+^ T cell functional states in human melanoma samples using a previously generated scRNA-Seq dataset (GSE120575). (**A)** Heatmap of select differentially expressed genes used to classify CD8^+^ T cells based on functional state: Memory/early activated (Mem/Early Activ.), Cytotoxic, Pre-dysfunctional (Pre-Dysfunc.), Dysfunctional (Dysfunc.), or Late Dysfunctional (Late Dysfunc.) states. (**B**) List of gene markers used to define each T cell state.

**Supplementary Fig. 8.**
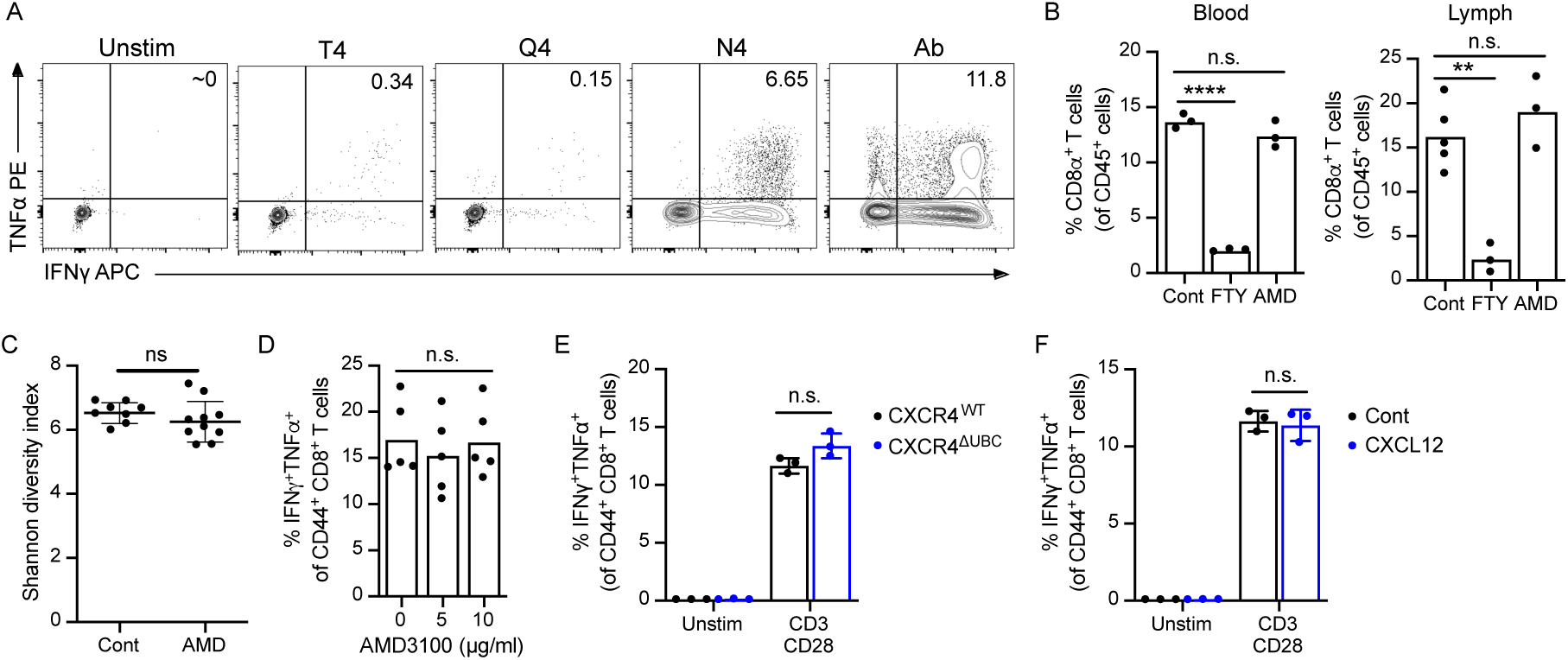
CXCR4 inhibition does not impact effector circulation or function. (**A**) Representative flow plots of IFN and TNF production by endogenous OVA-specific CD44^+^CD8^+^ T cells in the absence (Unstim) or presence of functional CD3/CD28 antibodies (Ab) or peptides: SIINFEKL (N4), SIIQFEKL (Q4), SIITFEKL (T4). (**B**) Frequency of CD8^+^ T cells in the blood (left) or lymph (right) following acute treatment with FTY720 (FTY) or AMD3100 (AMD). n=3 mice per group. (**C**) Shannon diversity index for TCR sequences detected in YUMMER1.7 tumors 7 days post treatment with AMD3100 (AMD; n=10) or vehicle control (Cont; n=8). (**D**) Frequency of IFN ^+^TNF ^+^ cells among endogenous CD44^+^CD8^+^ T cells following *ex vivo* stimulation with functional CD3/CD28 antibodies in the presence or absence of AMD3100. n=5 (**E**) Frequency of IFN ^+^TNF ^+^ cells among endogenous CXCR4^WT^ (n=3) or CXCR^ΔUBC^ (n=3) CD44^+^CD8^+^ T cells following *ex vivo* stimulation with functional CD3/CD28 antibodies. (**F**) Frequency of IFN ^+^TNF ^+^ cells among endogenous CD44^+^CD8^+^ T cells following *ex vivo* stimulation with functional CD3/CD28 antibodies in the presence (n=3) or absence (n=3) of recombinant murine 100 ng/ml CXCL12. For all graphs, each symbol represents one mouse; all error bars = standard deviation. One-way ANOVA (B and D), Mann-Whitney test (C), two-sided, unpaired student’s t-test (E and F). **p<0.01, n.s.=not significant.

